# Gut bacteria-derived serotonin promotes immune tolerance in early life

**DOI:** 10.1101/2022.09.25.509428

**Authors:** Katherine Z. Sanidad, Stephanie L. Rager, Hannah C. Carrow, Aparna Ananthanarayanan, Ryann Callaghan, Lucy R. Hart, Tingting Li, Purnima Ravisankar, Julia A. Brown, Mohammed Amir, Jenny C. Jin, Alexandria Rose Savage, Ryan Luo, Florencia Mardorsky Rowdo, M. Laura Martin, Randi B. Silver, Chun-Jun Guo, Jan Krumsiek, Naohiro Inohara, Melody Y. Zeng

## Abstract

The gut microbiome promotes immune system development in early life, but the neonatal gut metabolome remains undefined. Here, we demonstrate that, distinct from adults, the neonatal mouse gut is enriched with neurotransmitters, and specific bacteria produce serotonin directly while downregulating monoamine oxidase A to limit serotonin breakdown. Serotonin inhibits mTOR activation to promote regulatory T cells and suppress T cell responses both *ex vivo* and *in vivo* in the neonatal intestine. Oral gavage of serotonin into neonatal mice leads to long-term immune tolerance toward both dietary antigens and commensal bacteria as well as alterations of the gut microbiome. Together, our study has uncovered unique microbiome-dependent mechanisms to maximize serotonin in the neonatal gut and a novel role for intestinal serotonin to promote immune tolerance in early life.

## Introduction

Bacterial colonization of the gut in early life is a critical driver of the maturation of the gut and the development of the immune system (Backhed et al., 2015; Belkaid and Hand, 2014; Milani et al., 2017); this is mediated in part by the gut metabolome. Unlike the adult gut, the maturing neonatal gut is enriched with sugars and milk oligosaccharides, colonized by a dynamically changing gut microbiome, and inhabited by maturing immune cells under the influence of microbial signals. Recent studies have reported altered gut microbiome and metabolome in children with food allergies, asthma and neurodevelopmental defects (Fujimura et al., 2016; Levan et al., 2019; Sharon et al., 2019). Early life is also a crucial time window for the establishment of immune tolerance towards gut commensal bacteria as well as dietary and environmental antigens (Di Giovangiulio et al., 2015; Tsuji and Kosaka, 2008). It remains poorly understood whether/how the neonatal metabolome influences the development of immune tolerance in early life.

The gut has emerged as a major site of neurotransmitters, including dopamine and serotonin, which are produced mainly by epithelial enterochromaffin cells and exert both local and systemic effects on the enteric and central nervous systems (Sharon et al., 2016). Germ-free mice exhibit abnormal stress responses and fear extinction learning (Chu et al., 2019; Sudo et al., 2004; Wu et al., 2021) in part due to altered availability of metabolites or neurotransmitters for neuronal signaling in the absence of critical gut bacterial to facilitate this process. More evidence has emerged in recent years to shed light on gut bacteria as a critical regulator of neuroinflammation that underlies neurodevelopmental disorders (Bostick et al., 2022; Kim et al., 2017a; Needham et al., 2022). Our understanding of the regulation of neurotransmitters in the gut during early development, however, remains limited.

Serotonin (5-hydroxytryptamine or 5-HT), an essential neurotransmitter known to control gut motility, platelet function, and mood regulation, is also implicated in gut inflammatory diseases such as inflammatory bowel disease (IBD) or inflammatory bowel syndrome (IBS) (Di Giovangiulio et al., 2015); this is largely thought to be a consequence of indirect effects of 5-HT on enteric neurons. However, it remains unclear whether there is a direct crosstalk between 5-HT with gut immune cells. Up to date, our understanding of 5-HT synthesis in the gut is solely based on studies of adult animals; currently, the regulation and functions of 5-HT in the neonatal gut remain undefined.

In the present work, we report a neurotransmitter-enriched gut metabolome in the mouse neonatal small intestine. We further demonstrate that gut bacteria maximize 5-HT availability in the neonatal gut through 3 independent mechanisms: by directly producing 5-HT, inducing host tryptophan hydroxylase 1 to promote the conversion of tryptophan to 5-HT, and limiting the breakdown of 5-HT by downregulating monoamine oxidase A. Our findings suggest that 5-HT directly affects T cell metabolism to promote regulatory T cells while dampening helper T cell activation both in vitro and in vivo in the neonatal small intestine. Lastly, we show that exposure of 5-HT in the neonatal gut leads to long-term immune tolerance to both dietary antigens and gut commensal bacteria. Collectively, this work has uncovered a unique neurotransmitter-enriched metabolome in the neonatal gut and a novel gut metabolite-mediated mechanism to promote oral immune tolerance in early life.

## Results

### Enrichment of neurotransmitters in the neonatal intestine

The gut metabolome, collectively referring to the metabolites (small molecules) mainly produced by gut commensal bacteria, mediates the effects of gut bacteria on host cells. The microbiome and metabolome in the small intestine (SI), particularly in neonates, are not well characterized. We found distinct microbial populations in the small intestines of neonatal WT B6 mice (14 days postnatal; P14), with reduced diversity compared to that in adult mice (age > 8 wks) raised under specific pathogen-free (SPF) conditions (**Fig. 1A-D**). *Lactobacillus* species represent over 90% of bacteria in the neonatal SI, including *L. murinus*, which was more abundant in the neonatal small intestine (**Fig. 1C-D, Table S1**). Unbiased metabolomic profiling of SI luminal contents from neonatal (P14) and adult mice showed that the neonatal SI metabolome differed significantly from the adult SI metabolome (**Fig. 1E**). Of note, the metabolites that were enriched in the neonatal SI were almost exclusively neurotransmitters, including serotonin, hypotaurine, and acetylcholine (**Fig. 1F-G**). Consistently, KEGG pathways, such as serotonergic and cholinergic synapses, taurine and hypotaurine metabolism, and tryptophan metabolism, were upregulated in the neonatal SI, again highlighting a unique metabolic state in the neonatal SI (**Fig. 1H**). Together, our findings demonstrate unique gut microbiome and metabolome landscapes in the neonatal SI with distinct enrichment of neurotransmitters.

**Fig. 1.**
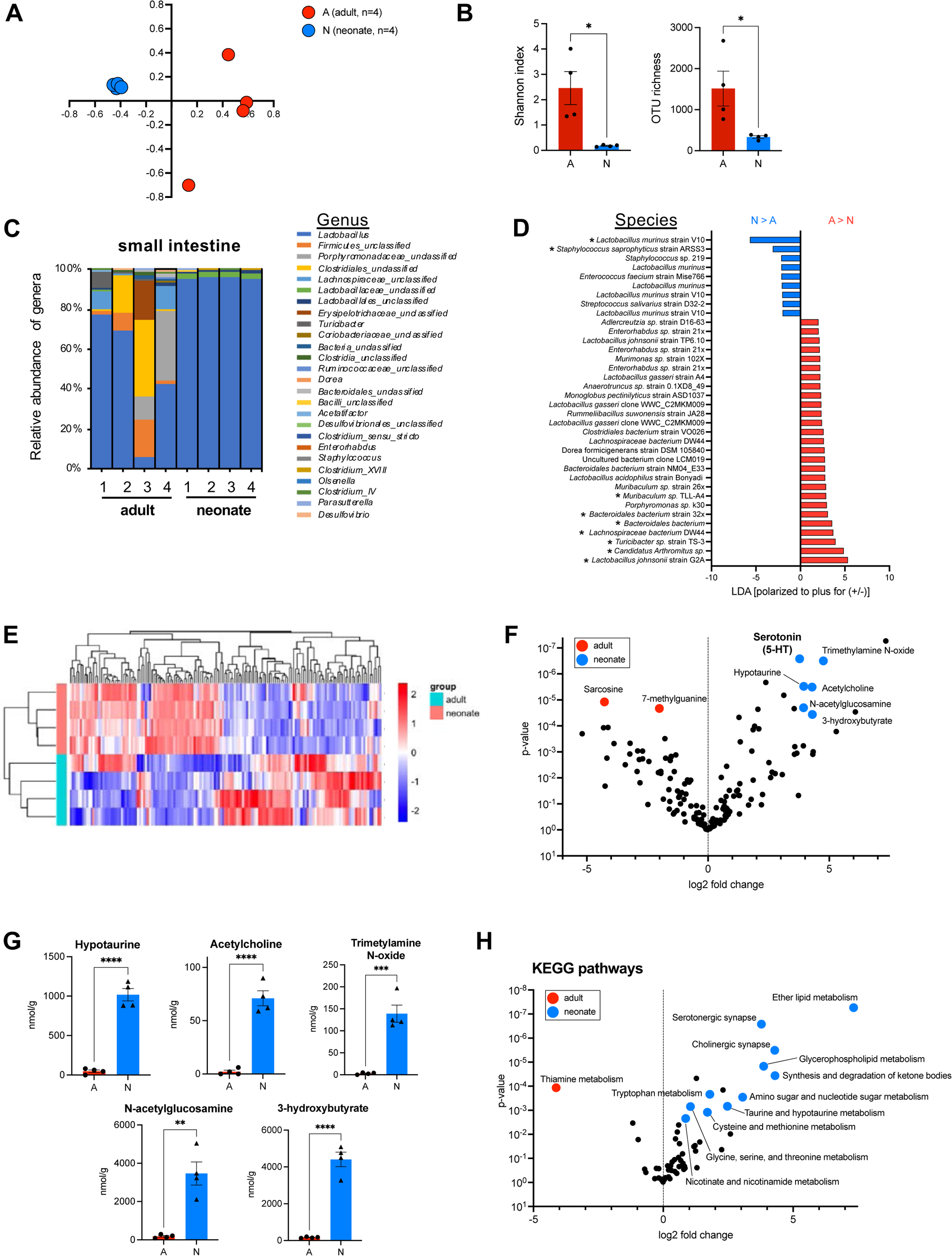
Enrichment of neurotransmitters in the neonatal intestine. (**A**) β-diversity represented by NMDS analysis and (**B**) α-diversity measured by Shannon index and OTU richness of gut microbiomes in SI of SPF adult (>8 weeks) and neonatal (2-week-old) mice. (**C-D**) 16S rRNA sequencing analysis of (C) relative abundance of genera and (D) LDA of specific OTUs (labels represent bacteria with highest homology; * = abundance > 0.5%) found in SI of SPF adult and neonatal mice. (**E**) Heatmap of ∼ 500 metabolites in SI luminal contents of SPF adult and neonatal mice through high-throughput metabolomics analyses. (**F**) Volcano plot representing fold change of metabolites abundant in SI luminal contents of adult and neonatal mice. (**G**) Concentrations of specific metabolites from high throughput metabolomics data in adult and neonatal SI. (**H**) Volcano plot representing fold change of KEGG pathways associated with SI luminal metabolites between adult or neonatal mice. \**P*<0.05, \*\*\**P*<0.001, \*\*\*\**P*<0.0001. Adult mice age >8 weeks, neonatal mice age=P14. See also **Table S1**.

### Serotonin (5-HT) biosynthesis in the neonatal gut is driven by the gut microbiome

A previous study has shown that 5-HT biosynthesis in adult mice is promoted by selective gut bacteria via upregulation of tryptophan hydroxylase 1 (TPH1), which converts tryptophan to 5-HT, in enterochromaffin cells (Yano et al., 2015b). The regulation of 5-HT biosynthesis in the neonatal gut, however, remains unexplored. We found elevated levels of 5-HT in the neonatal SI compared to the adult SI, despite similar levels of tryptophan, the precursor of 5-HT (**Fig. 2A**). The metabolized product of 5-HT, 5-hydroxyindoleacetic acid (5HIAA), however, was almost undetectable in the neonatal SI (**Fig. 2A**), suggesting reduced breakdown of 5-HT. Interestingly, we found higher levels of 5-HT in the SI luminal contents and tissues from SPF WT P14 neonates compared to that in GF WT neonates, and to a lesser extent in the colon luminal contents (**Fig. 2B-D, S1A**). In contrast, similar levels of 5-HT were detected in SPF and GF adult luminal contents (**Fig. 2B**). These findings suggest a more profound role for the gut microbiome in 5-HT biosynthesis in the neonatal gut than in the adult gut. In addition, similar levels of 5-HT were detected in colon tissues, plasma, and brains between neonates and adults, in both SPF and GF conditions, suggesting neither age nor gut microbiome is a factor in 5-HT biosynthesis in these tissues (**Fig. S1A-B**). Together, our findings elucidate a critical role for the gut microbiome in the regulation of SI 5-HT biosynthesis during early life.

**Figure 2.**
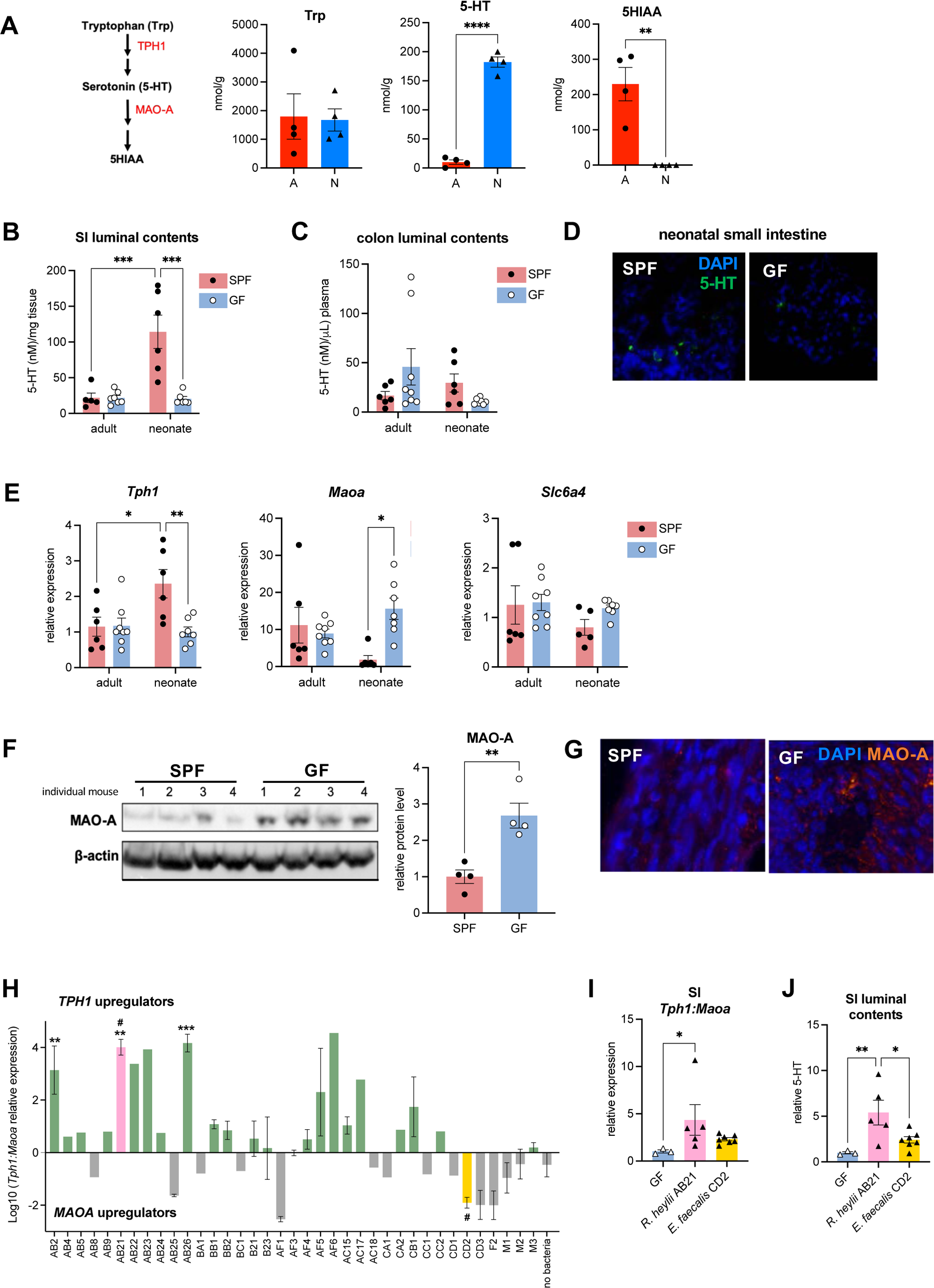
Serotonin (5-HT) biosynthesis in the neonatal intestine is driven by the gut microbiome. (**A**) 5-HT synthesis pathway in the gut and concentrations of Tryptophan (Trp), 5-HT, and 5HIAA in adult and neonatal SI. (**B-C**) Concentrations of 5-HT in SPF and GF adult (> 8-week-old) and neonatal (P14) mice measured by LCMS in luminal contents of (**B**) SI and (**C**) colon. (**D**) Immunofluorescence staining of 5-HT in the small intestine of SPF and GF neonatal mice. (**E**) RT-qPCR analysis of *Tph1*, *Maoa*, and *Slc6a4* gene expression in SI of SPF/GF adult and neonatal mice. (**F**) Western blot analysis and (**G**) immunofluorescence staining of MAO-A in SI of SPF and GF neonatal mice. (**H**) Ratio of *TPH1:MAOA* expression in HT-29 colon cells treated with supernatants of bacterial isolates from neonatal mouse SI. Significance is shown compared to no bacteria control (cells not treated with bacterial supernatant); # designates isolates used for in vivo experiments. (**I-J**) (**I**) RT-qPCR analysis of the ratio of *Tph1:Maoa* expression in SI of P16 GF neonatal mice monocolonized with designated isolates in **Fig. 2H**. (**J**) 5-HT concentrations in SI luminal contents of P16 mice monocolonized with designated isolates in **Fig. 2H** measured by ELISA. \**P*<0.05, \*\**P*<0.01, \*\*\**P*<0.001, \*\*\*\**P*<0.0001. Adult mice age >8 weeks, neonatal mice age=P14. See also **Figure S1** and **Table S2**.

### The neonatal gut microbiome inhibits MAO-A expression to reduce 5-HT breakdown

To further ascertain the mechanisms by which the gut microbiome regulates 5-HT biosynthesis in the neonatal intestine, we found the expression of TPH1, the rate-limiting enzyme involved in synthesizing 5-HT from tryptophan, was ∼2.5-fold higher in the SI of SPF WT neonates compared to that in the SI of GF WT neonates (**Fig. 2E**). More importantly, monoamine oxidase A (MAO-A), which converts 5-HT into 5HIAA, was expressed at significantly lower (∼15 fold) levels in the SI of SPF WT neonates (**Fig. 2E**), consistent with the absence of 5HIAA in SPF neonatal SI (**Fig. 2A**). Moreover, at the protein level, MAO-A was substantially decreased in the SI of SPF neonates compared to in GF neonates (**Fig. 2F-G**), underscoring a crucial role for the gut microbiome in the downregulation of MAO-A in the neonatal SI. In contrast, the gut microbiome appeared dispensable for the expression of *Tph1* and *Maoa* in the neonatal colon (**Fig. S1C**), and the SI and colon expression of *Slc6a4*, the gene that encodes the serotonin reuptake transporter (SERT) (**Fig. 2E, S1C**). Human colon organoids treated with SI luminal contents from SPF neonatal mice induced a higher *TPH1:MAOA* expression ratio than in organoids treated with GF SI luminal contents (**Fig. S1D)**, consistently suggesting a direct role for neonatal SI gut bacteria in the modulation of *Tph1* and *Maoa* expression in the neonatal SI. Altogether, our findings demonstrate the neonatal gut microbiome promotes 5-HT in the neonatal SI in part by increasing TPH1 expression and reducing MAO-A expression to maximize the availability of intestinal 5-HT during the early developmental stage.

As proof of concept, we screened bacterial isolates from the P14 WT B6 mouse SI for modulation of *TPH1* and *MAOA* expression in human colonic epithelial HT-29 cells. A few bacterial isolates, including AB21, which was identified as *Rodentibacter heylii* (**Fig. 2H)**, induced the highest *TPH1:MAOA* ratio, referred to as TPH1 upregulators (**Fig. 2H; Table S2**). GF WT B6 neonates were successfully mono-colonized with AB21 indirectly by orally inoculating the dam at P3 (**Fig. S1E**). At P14, these neonates exhibited a higher *Tph1:Maoa* ratio and increased amounts of luminal 5-HT in the SI, compared to neonates that were similarly mono-colonized with a MAO-A upregulating bacterial isolate (CD2; *E. faecalis*) (**Fig. 2H-J, S1F-G**). These findings demonstrate that bacteria from the neonatal SI influence 5-HT intestinal availability in part by modulating the expression of TPH1 and MAOA.

### Gut bacteria are the major 5-HT producers in the neonatal intestine

Enterochromaffin (EC) cells are the major 5-HT producers in the adult intestine (Lund et al., 2018). However, the source of 5-HT in the neonatal intestine remains unclear. We detected comparable 5-HT levels by ELISA and immunofluorescence staining for SI 5-HT in epithelial cell-specific *Tph1*-deficient *Tph1 ^fl/fl^ x Villin Cre* and *Tph1 ^fl/fl^* littermate P14 mice (**Fig. 3A-C, S2A**), suggesting EC cells are not major 5-HT producers in the neonatal intestine, in stark contrast to the adult intestine. Hematopoietic cells have been shown to produce 5-HT, particularly mast cells (Herr et al., 2017; Kushnir-Sukhov et al., 2007). Similar numbers of lamina propria mast cells (CD117+FcεRI+) were found the SI of SPF and GF WT P14 neonates (**Fig. S2B**). Conversely, immunofluorescence co-staining for 5-HT and avidin, a mast cell marker, showed similar numbers of 5-HT+ mast cells in the SI of SPF and GF WT P14 neonates (**Fig. S2C**), as well as in the SI of P14 *Tph1 ^fl/fl^ x Villin Cre* and heterozygous littermate mice (**Fig. S2D-E**). Together, our findings suggest that, distinct from the adult intestine, enterochromaffin cells or mast cells are not a major source of 5-HT in the neonatal intestine.

**Figure 3.**
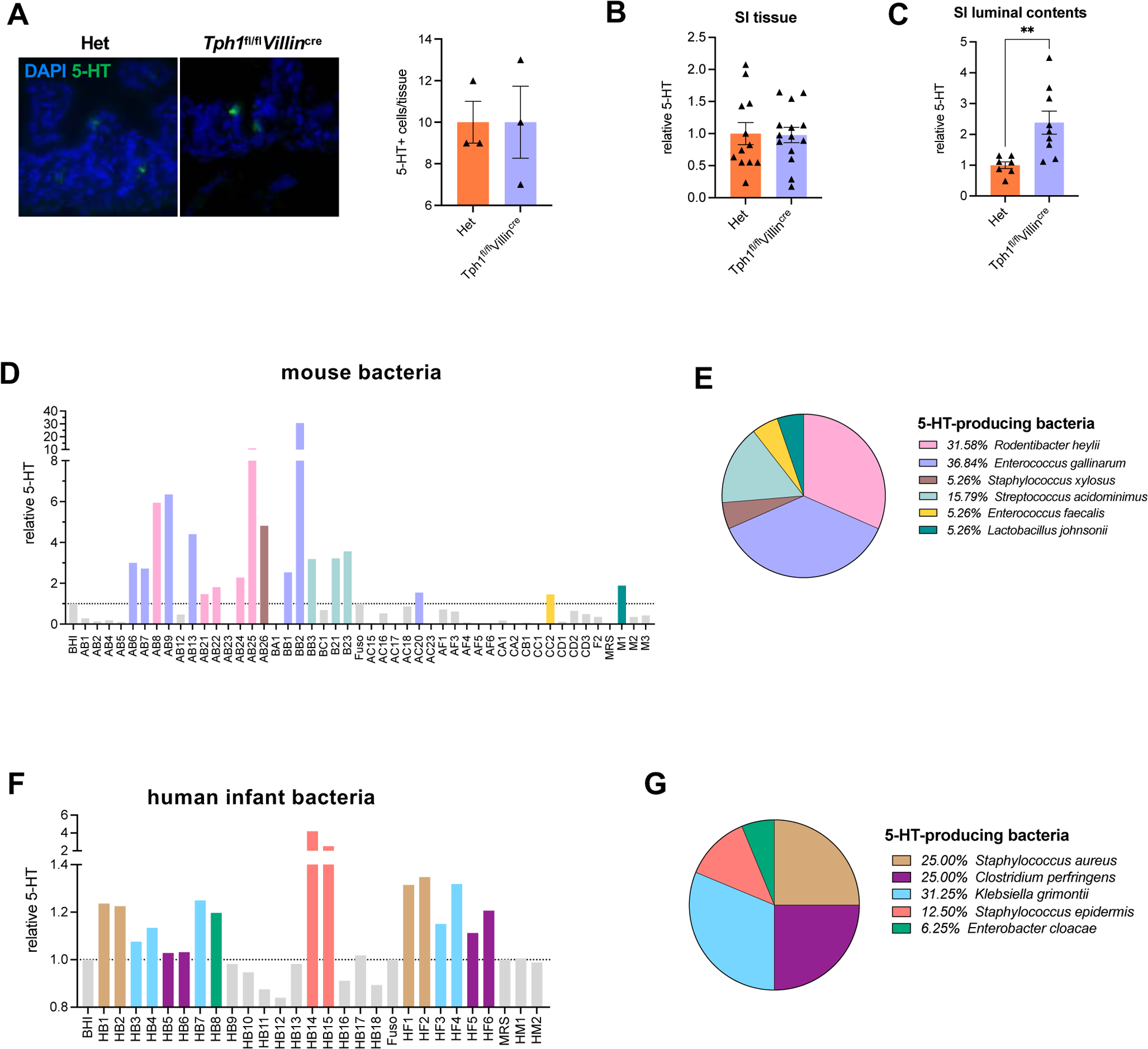
Gut bacteria are the major 5-HT producers in the neonatal intestine. (**A**) Representative images of immunofluorescence staining of 5-HT in SI of *Tph1*^fl/fl^*Villin^cre^* neonatal (2-week-old) mice or heterozygous (Het) littermates and quantification of 5-HT stained cells/tissue. (**B-C**) ELISA analysis of 5-HT concentrations in (**B**) SI tissue and (**C**) luminal contents of *Tph1*^fl/fl^*Villin*^cre^ neonatal mice or Het littermates. (**D**) ELISA analysis of 5-HT in supernatants of bacteria isolated from SI of neonatal mice. (**E**) Percentage of bacterial isolates from SI of neonatal mice capable of producing 5-HT. (**F**) ELISA analysis of 5-HT in supernatants of bacteria isolated from stool of term healthy infants without antibiotics. (**G**) Percentage of bacterial isolates from stool of term healthy infants without antibiotics capable of producing 5-HT. \*\**P*<0.01. Neonatal mice age = 2 weeks. See also **Figure S2** and **Table S3**.

Bacteria such as *Escherichia coli* have been shown previously to metabolize tryptophan and generate 5-HT (Nzakizwanayo et al., 2015). To investigate the possible existence of 5-HT-producing bacteria in the neonatal SI, we isolated bacteria from P14 SPF WT neonates aerobically and anaerobically and generated a library of >30 bacterial isolates. Normalized bacterial supernatants were measured for 5-HT by ELISA; almost half of the bacterial isolates produced 5-HT (**Fig. 3D**). Close to 1/3 of 5-HT-producing bacterial isolates were *Rodentibacter heylii,* and another 1/3 were *Enterococcus gallinarum* (**Fig. 3E; Table S3**). In addition, using a similar approach, we showed that 16 out of 26 bacterial isolates from healthy human infant (gestational age > 37 weeks) stool specimens during the first 2 weeks of life produced 5-HT, which were predominantly *Staphylococcus aureus*, *Clostridium perfringens*, *Klebsiella grimontii*, *Staphylococcus epidermis*, and *Enterobacter cloacae* (**Fig. 3F-G**). Of note, our data only reflect the 5-HT-producing bacteria that were culturable in the laboratory setting; there are likely other 5-HT-producing bacteria that are not culturable. Collectively, our findings demonstrate the microbiome as the major source of 5-HT in the neonatal gut.

### 5-HT regulates T cell response *in vitro* and *in vivo* in the neonatal SI

5-HT receptors or 5-HT re-uptake transporters (SERTs) are expressed in a variety of cell types, including both hematopoietic and non-hematopoietic cells (Herr et al., 2017). However, most clinical or animal studies of 5-HT were performed in adults; the physiologic significance of gut-derived 5-HT during early life remains poorly understood. 5-HT was previously shown to promote the expansion of regulatory T cells (Tregs) in the central nervous system (CNS) (Ito et al., 2019; Sacramento et al., 2018). It remains unclear if intestinal 5-HT directly affects T cell response in the gut. We found a selective reduction in lamina propria (LP) Tregs in the SI of P14 GF mice compared to SPF neonates (**Fig. 4A**), suggesting a role for the gut microbiome in the development of Tregs in the SI of neonates. Gut microbiome-derived short chain fatty acids (SCFAs) such as butyrate and propionate, which are well described to induce Tregs in the colon (Smith et al., 2013), were almost absent in the neonatal SI compared to adult SI (**Fig. 4B**). This suggests that the gut microbiome promotes Treg development in the neonatal SI by a SCFA-independent mechanism. We found that 5-HT directly influences T cell response in vitro, as incubation of splenic T cells with 5-HT for 48 h *in vitro* increased Tregs while reducing the production of IFN-γ and IL-17A production in both CD4+ T cells (**Fig. 4C, S3A**) and CD8+ T cells (**Fig. S3B**).

**Figure 4.**
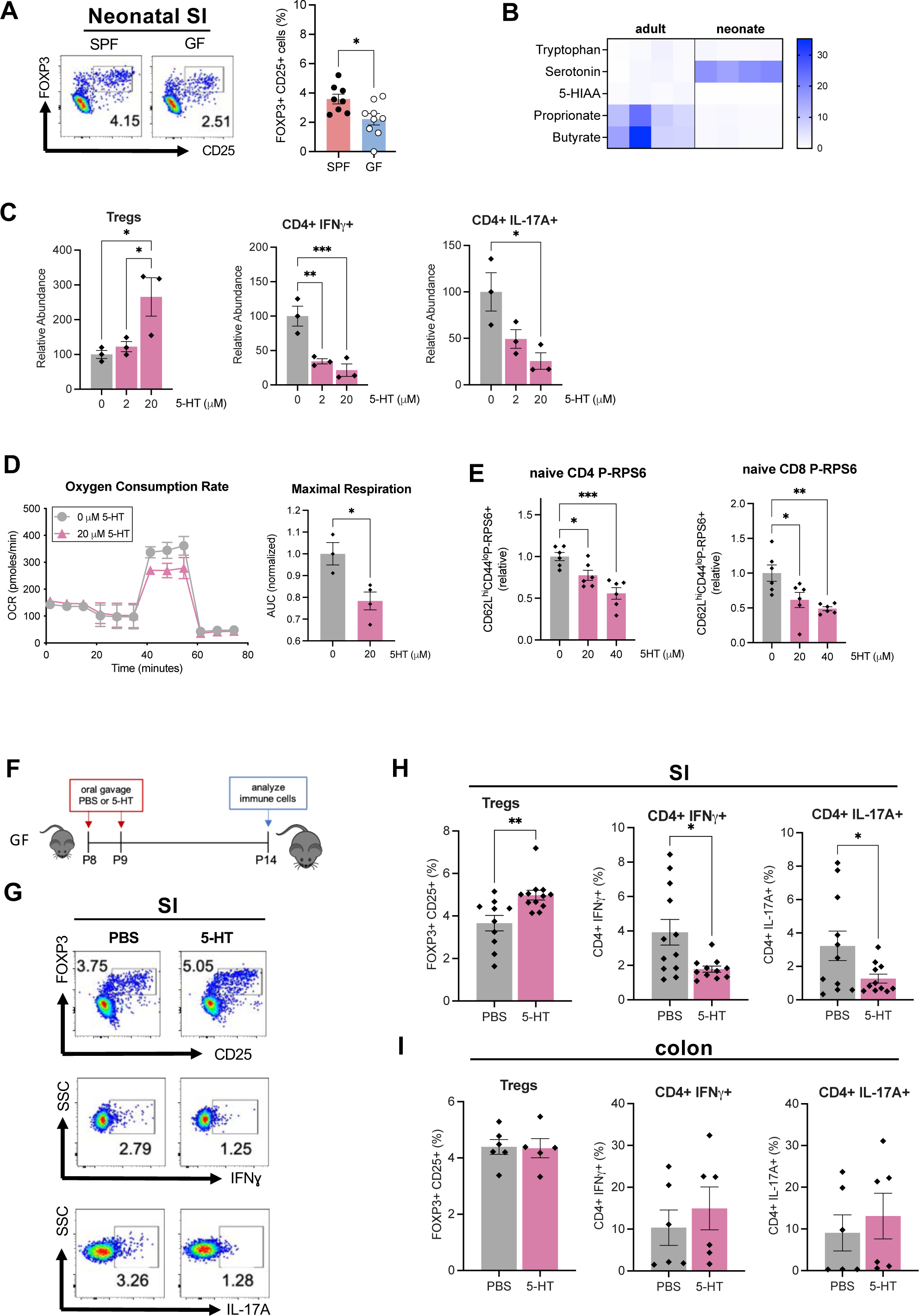
5-HT promotes Treg development in vitro and in vivo in the neonatal SI. (**A**) Representative flow plots (top) and flow cytometry analysis (bottom) of Tregs isolated from the lamina propria of SI and colon of SPF and GF neonatal (2-week-old) mice. (**B**) Heatmap of relative metabolite concentrations in SI of SPF adult (>8 weeks old) and neonatal (2-week-old) mice. (**C**) Flow cytometry analysis of mouse splenic CD4+ T cells stimulated with 5-HT for 48 h *ex vivo*. (**D**) Seahorse real-time cell metabolic analysis of oxygen consumption rate (left) and maximal respiration (right; AUC calculated between 40-60 min) of mouse splenic T cells stimulated with 5-HT for 3 h *in vitro.* (**E**) Flow cytometry analysis of mouse naïve CD3+ P-RPS6+ splenic T cells stimulated with 5-HT for 24 h. (**F**) Schematic of experiment of GF neonatal mice orally treated with PBS or 5-HT. (**G-H**) (**G**) Representative flow plots and (**H**) flow cytometry analysis of lamina propria cells in the SI of GF neonatal mice orally gavaged with PBS or 5-HT. (**I**) Flow cytometry analysis of lamina propria cells in the colon of GF neonatal mice orally gavaged with PBS or 5-HT. \**P*<0.05, \*\**P*<0.01, *** *P*<0.001. Neonatal mice age = 2 weeks. See also **Figure S3**.

Nutrients control cell fate commitment and responses of different T cell subsets independent of antigens presented by antigen-presenting cells and cytokines (Chapman and Chi, 2014). Tryptophan is known to affect T cell effector functions via the kynurenine pathway, in which tryptophan is converted into kynurenine by indoleamine 2,3-dioxygenase (IDO) (Wei et al., 2017); however, it remains unknown whether the serotonin pathway can independently impact T cell metabolism. Through a Seahorse assay, we found that incubation with 5-HT reduced oxygen consumption by mouse splenic T cells, suggesting 5-HT directly changes T cell metabolism (**Fig. 4D**). Mammalian target of rapamycin (mTOR) signaling is critical for T cells to integrate immune signals and metabolic cues for their proper maintenance and activation. mTOR activation promotes the differentiation of Th1, Th2, and Th17 effector cells while inhibiting the development of Tregs (Delgoffe et al., 2009). We found that splenic naïve CD4+ or CD8+ T cells incubated with 5-HT for 24 h had lower levels of phosphorylated-ribosomal protein S6 (P-RPS6) (**Fig. 4E, S3C-D**), a biomarker of mTOR pathway activation (Ma and Blenis, 2009). This suggests that 5-HT alters the metabolic state of T cells in part via inhibition of the mTOR pathway, thus leading to Treg expansion while diminishing T cell activation.

Furthermore, based on the 5-HT concentrations in the SI luminal contents of SPF neonates determined by LC-MS analysis (**Fig. 2B)**, we orally gavaged 7.8 μg 5-HT into GF neonates at P8 and P9, and isolated LP cells from the SI and colon at P14 (**Fig. 4F**). Consistently, we found increased numbers of Tregs and reduced numbers of IL-17A+ or IFNγ+ T cells in the SI, but not the colon of 5-HT-treated GF neonates (**Fig. 4G-I, S3E**). Collectively, these studies demonstrate that 5-HT promotes the development of Tregs and dampens inflammatory IL-17A+ and IFNγ+ T cells both *in vitro* and in the neonatal SI *in vivo*.

### Exposure to intestinal 5-HT in early life contributes to long-term oral tolerance to dietary antigens

Oral immune tolerance to dietary antigens is established in early life, which is facilitated in part by intestinal Tregs (Pabst and Mowat, 2012). To determine if 5-HT contributes to oral immune tolerance toward dietary antigens in neonates, GF neonates were orally gavaged with 5-HT at P8 and P9, sensitized through oral gavage of OVA to the dam at P12, P15, P18, and P21, challenged with OVA subcutaneously at 5 weeks of age, re-challenged at 7 weeks of age, and finally sacrificed 3 days later (**Fig. 5A**). 5-HT-treated mice had lower plasma anti-OVA IgG and anti-OVA IgE antibodies compared to that in the PBS-treated mice (both sensitized and challenged similarly) as well as the control mice that were neither treated with 5-HT nor sensitized with OVA in early life (**Fig. 5B**). There was a similar trend for lower plasma OVA-specific IgM antibodies in 5-HT-treated mice (**Fig. S4A**). We found a decrease in splenic IFN-γ and IL-4-producing CD4+ T cells, macrophages, and neutrophils (**Fig. 5C, S4B**), as well as a significant reduction of B cells in the mesenteric lymph nodes (mLN) in 5-HT-treated mice challenged with OVA (**Fig. S4C**). Taken together, these findings suggest that exposure to intestinal 5-HT during early life promotes long-term systemic immune tolerance to oral antigens.

**Figure 5.**
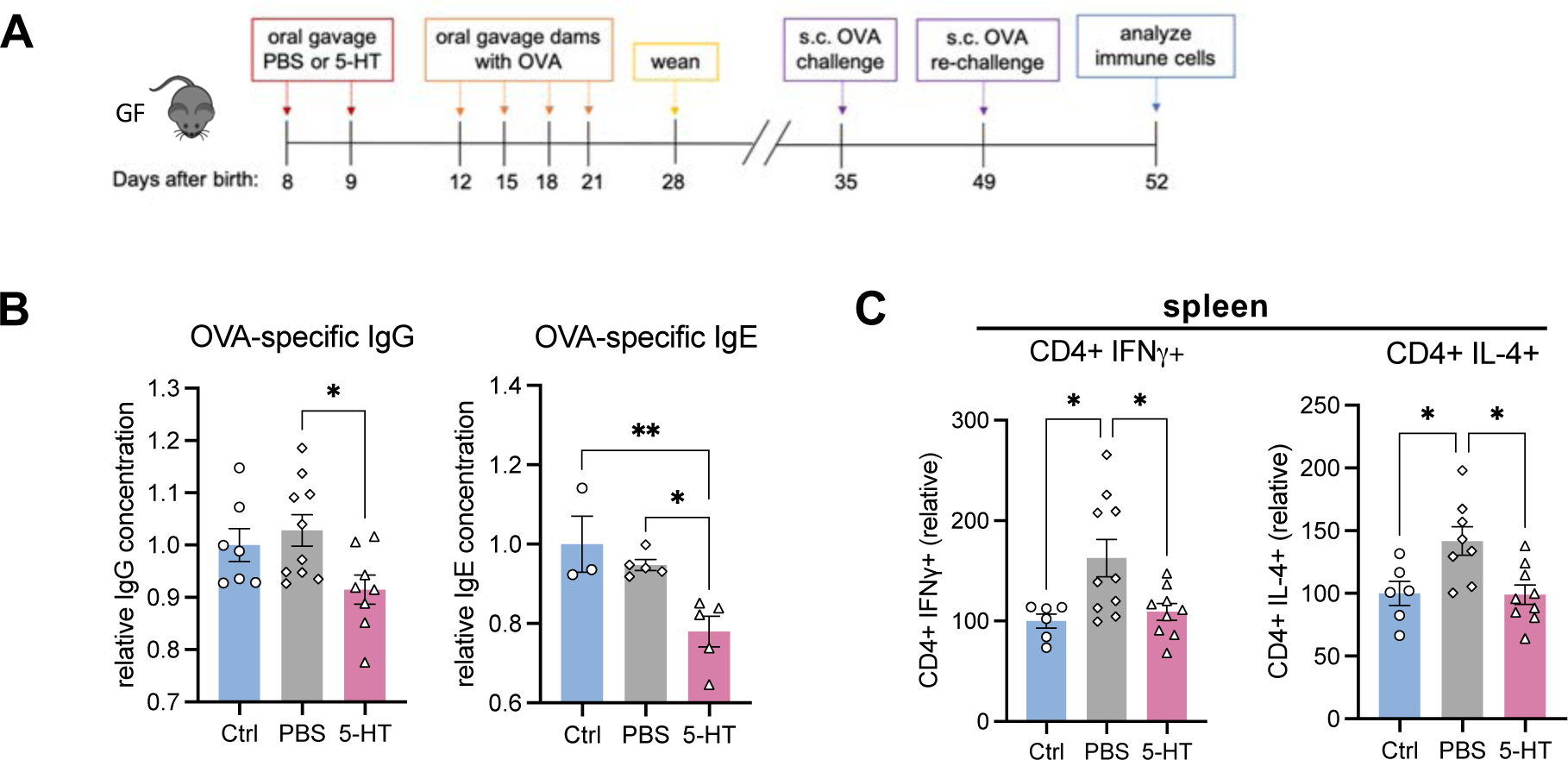
5-HT in the neonatal gut promotes long-term immune tolerance to oral antigens. (**A**) Schematic of mouse model of oral tolerance to OVA. (**B**) ELISA analysis of OVA-Specific IgG and IgE in plasma and (**C**) flow cytometry analysis of immune cells in spleen of mice used in model of oral tolerance to OVA.. \**P*<0.05, \**P*<0.01. See also **Figure S4**.

### Neonatal intestinal 5-HT promotes immune tolerance to gut commensal bacteria

Tregs in the SI suppress immune responses toward gut commensal bacteria (Di Giovangiulio et al., 2015; Sefik et al., 2015; Wawrzyniak et al., 2017). To test if 5-HT could also influence tolerance to commensal gut bacteria, 5-HT was orally gavaged to GF WT neonatal mice from P8-9 (**Fig. 6A**). At P14, the dam was gavaged with SI and colon luminal contents from SPF WT B6 P14 neonates to allow colonization of gut commensal bacteria and transfer of these bacteria to the PBS or 5-HT-treated littermates that were housed together with the dam. Two weeks later, comparable bacterial colonization in both groups of littermates was confirmed (**Fig. S5A**). Compared to GF mice and PBS-treated littermates, 5-HT-treated littermates had a higher abundance of Tregs and lower abundance of Th17 cells in the colons (**Fig. 6B**), as well as fewer Th17 cells in the mLN (**Fig. 6C**). Additionally, we found lower abundances of other immune cells including cytokine-producing-T cells, B cells, and dendritic cells in the intestines (**Fig. S5B-C**) and mLNs (**Figs. 6C, S5D**) of 5-HT-treated neonatal mice compared to the GF and PBS-treated-neonatal mice.

**Figure 6.**
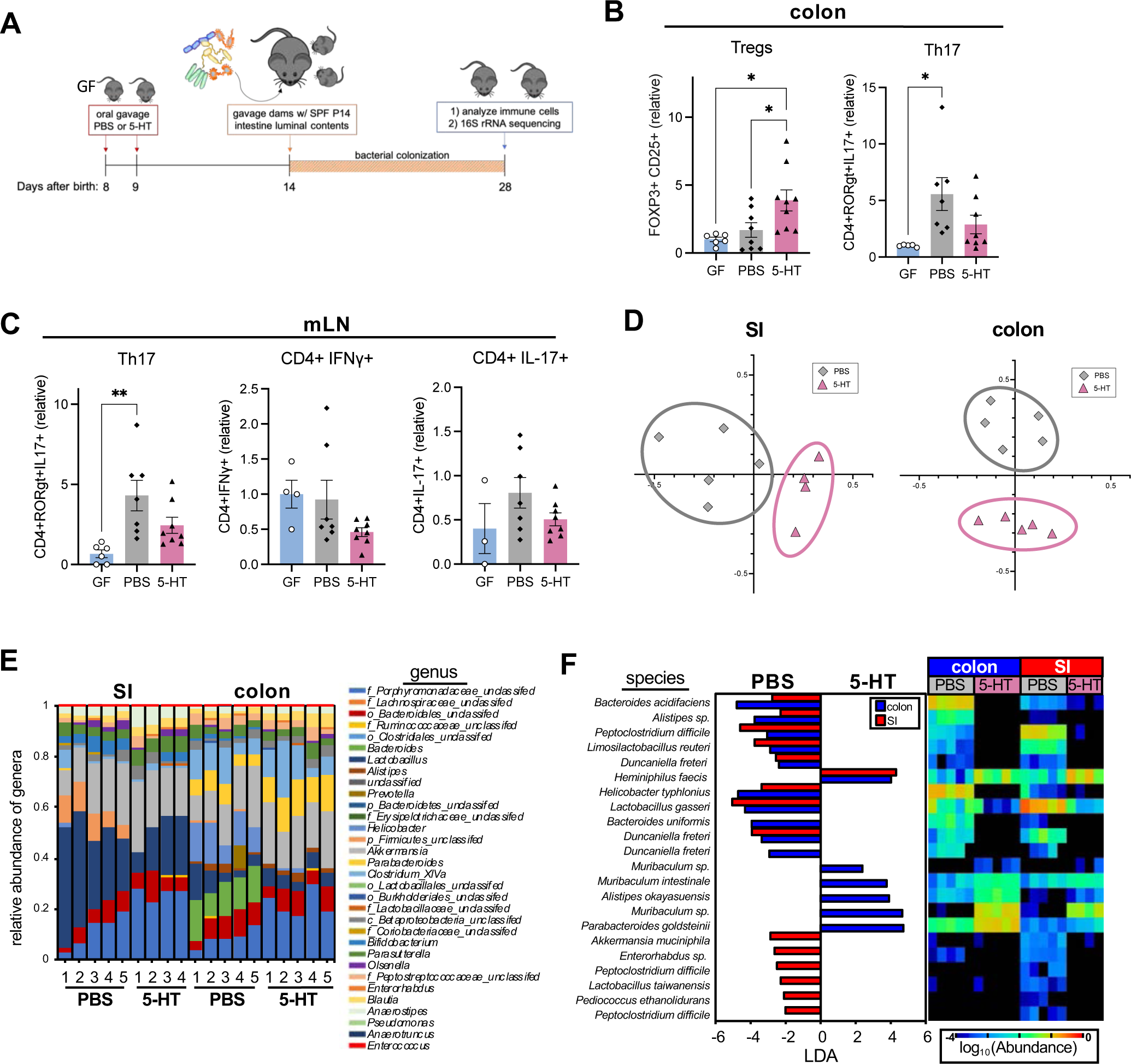
5-HT in the neonatal gut promotes immune tolerance to gut commensal bacteria. (**A**) Schematic of the experimental design to test the role of 5-HT in immune tolerance to commensal bacteria. GF neonates from the same litter were gavaged with 5-HT or PBS at P8 and P9 and remained co-housed through P28. At P14, the biological dam was orally gavaged with gut luminal and fecal bacteria from P14 SPF WT neonates (to allow transfer of bacteria to the offspring). All the mice were housed in the same cage. At P28, gut immune cells and microbiome were analyzed. (**B-C**) Flow cytometry analysis of immune cells from (**B**) laminal propria of the colon and (**C**) mLN of GF mice given 5-HT or PBS and colonized with commensal bacteria. (**D**) NMDS analysis of 16S rRNA sequencing data of gut microbiota of mice treated with PBS or 5-HT. (**E-F**) 16S rRNA sequencing data of (**E**) relative abundance of genera and (**F**) LDA and relative abundance of specific bacteria (labels represent bacteria with highest homology) in SI and colon of mice given 5-HT or PBS. \**P*<0.05, \**P*<0.01. See also **Figure S5**.

Furthermore, to determine if altered immune responses due to 5-HT exposure would affect the gut microbiome, through 16S rRNA sequencing at P28, we found distinct gut microbiomes in both the SI and colon between the PBS-and 5-HT-treated littermates that acquired the gut bacteria from the same biological dam at P14 (**Fig. 6D-F**). At the species level, *Muribaculum sp*., including *Muribaculum intestinale*, and *Parabacteroides goldsteini* were more abundant in the colons of 5-HT-treated offspring, which had lower abundance of *Bacteroides acidifaciens* and *Bacteroides uniformis* (**Fig. 6F**). Increased abundance of *Heminiphilus faecis* was found in both the colons and SIs of 5-HT-treated offspring (**Fig. 6F**). Since the two small doses of 5-HT were given to the neonates at P8-9, which was 6 days prior to maternal colonization, the changes in the gut microbiome in 5-HT offspring were likely attributed to altered immune cell responses induced by 5-HT in the offspring. Together, these findings suggest intestinal 5-HT influences gut microbe-immune cell interactions by promoting Tregs and suppressing Th17 cells, and as a result, both the immune landscape and the gut microbiome in the neonatal gut are altered after exposure to 5-HT.

## Discussion

As a major site of nutrient absorption, the small intestine also harbors a diverse gut microbiome that thrives on the unique nutrients available in the small intestine, including sugars, fats, minerals and proteins. Despite growing literature characterizing the fecal microbiome and its role in health and disease, the microbiome of the human small intestine remains understudied due to the difficulty of sampling small intestinal contents (Mittal et al., 2017). The small intestine microbiome in early life is even far less understood. Our novel finding of unique enrichment of key neurotransmitters in the mouse neonatal small intestine underscores the need for independent efforts to interrogate and understand the neonatal microbiome and metabolome. Studies in recent years have provided compelling evidence for the gut as a major site for neurotransmitter generation, which are mainly produced by specialized epithelial EC cells (Sharon et al., 2016), based on studies of almost exclusively adult animals. The enriched neurotransmitters in the neonatal gut may be critical to support the demand for these key neurotransmitters during the early developmental period, both within the gut and beyond. The evolutionary benefits of increased abundance of neurotransmitters in the neonatal gut remain to be investigated. Gut-derived neurotransmitters typically are unable to cross the blood-brain barrier but can be circulated systemically via various mechanisms, such as the extra-intestinal transport of serotonin via platelets. The enteric nervous system (ENS) orchestrates gastrointestinal behavior independently of the CNS (Rao and Gershon, 2016). The development of ENS may be influenced by the increased availability of neurotransmitters in the neonatal small intestine, as ENS neurons form during the early postnatal period and continue through P21 in mice. Understanding the evolutionary benefits of enhanced neurotransmitter availability in the neonatal gut may shed light on GI disorders that stem from an imbalance in one of the key neurotransmitters during early life.

Our studies indicate a crucial role for the gut microbiome in the increased abundance of 5-HT in the neonatal SI via three major independent mechanisms – direct production of 5-HT, induction of TPH1 (∼2.5 fold higher than in germ-free SI), and downregulation of MAO-A to inhibit 5-HT breakdown. In contrast, 5-HT in the adult intestine is produced mainly by EC cells via TPH1 and the expression of MAO-A is far higher than that in the neonatal gut, the latter likely contributing to diminished levels of 5-HT in the adult gut. Of note, in our studies, we found that the gut microbiome is dispensable for 5-HT synthesis in both the SI and colon in adult mice, which departs from a prior study reporting that the gut microbiome induces a modest 1.5-fold higher expression of TPH1 in the adult colons compared to that in GF adult colons (Yano et al., 2015a). Thus, our findings suggest a more profound role for the gut microbiome in promoting 5-HT in the neonatal gut. This also raises the question of how perturbations of the gut microbiome during early life, such as due to antibiotic use, may impact gut 5-HT availability. This is an important question that requires future investigation.

Previous studies reported that 5-HT reduces IFN-γ and IL-17 production and promotes Tregs in patients with multiple sclerosis (MS) and in animal models of MS (Rao and Gershon, 2016; Wan et al., 2020), suggesting a direct role for 5-HT to modulate T cell response in the CNS. Whether this immunomodulatory function of 5-HT extends to the neonatal gut, however, remains unexplored. We found that oral gavage of 5-HT to GF neonates promotes Tregs and suppresses IFN-γ+ or IL-17A+ T cells. Mechanistically, 5-HT treatment changes the metabolic state of T cells, evidenced by a reduction in mTOR activation and oxygen consumption in 5-HT-treated T cells. Inhibition of mTOR activation leads to differentiation of Foxp3+ Tregs and suppression of Th1, Th2 or Th17 differentiation (Chapman et al., 2020; Chapman and Chi, 2014). Oral immune tolerance established in early life, in part mediated by gut Tregs, is critical to maintain local and systemic immune unresponsiveness to dietary antigens. The long-term effect of 5-HT in promoting immune tolerance to oral antigens, such as OVA in our study, demonstrates an example of the evolutionary benefits of increased availability of gut 5-HT during early life that is facilitated by unique bacteria in the neonatal gut. Furthermore, we also demonstrate that by shaping the immune landscape in the neonatal gut, exposure to 5-HT leads to significant changes in the gut microbiome. 5-HT-induced alterations of the neonatal gut microbiome may independently shape the development of neonatal gut immune cells but will require further investigation. Collectively, our studies have demonstrated a unique neurotransmitter-enriched metabolome in the neonatal gut and unraveled novel immune functions of 5-HT in the neonatal gut to promote immune tolerance (proposed model in **Fig. S6**). Our findings might have implications for a potential link between imbalanced neurotransmitters, including 5-HT, in the neonatal gut during early life and the development of GI inflammatory disorders and systemic autoimmunity in later life.

## Acknowledgments

We thank G.F. Sonnenberg for critical review of the manuscript, R. Pinefor and S. Paisner from the Weill Cornell Gnotobiotic Animal Facility for assistance with gnotobiotic animal studies, and T. Miller for assistance with flow cytometry.

## Funding

National Institute of Health grant R01HD110118 (MYZ)

National Institute of Health grant R01HL169989 (MYZ)

National Institute of Health grant R21CA270998 (MYZ)

National Institute of Health grant K01DK114376 (MYZ)

Hartwell Foundation Individual Biomedical Research Award (MYZ)

The Starr Cancer Consortium (MYZ)

Gale and Ira Drukier Institute for Children’s Health at Weill Cornell Medicine (MYZ)

Children’s Health Council at Weill Cornell Medicine (MYZ)

Center for Immunology and Office of Academic Integration of Cornell University (MYZ)

Center for IBD Research at Weill Cornell Medicine (MYZ)

National Center for Advancing Translational Sciences (NCATS) grant 2TL1TR2386 of the Clinical and Translational Science Center at Weill Cornell Medicine (KZS and JAB)

Biocodex Microbiota Foundation (JAB)

Hartwell Foundation Postdoc Fellowship (KZS)

National Institute of Health grant 1DP2HD101401 (CJG)

National Institute of Health grant DK135816 (CJG)

National Institute of Health grant AI172027 (CJG)

National Institute of Health grant DK132244 (CJG)

The Kenneth Rainin Foundation (CJG)

W.M. Keck Foundation (CJG)

## Author contributions

Conceptualization: MYZ Methodology: KZS, MYZ

Investigation: KZS, SLR, HCC, AA, RC, LRH, TL, PR, JAB, MA, JLJ, ARS, RL, FMR, MLM, RBS, JMP, CJG, NI

Visualization: JK, NI, KZS

Funding acquisition: MYZ

Project administration: MYZ

Supervision: MYZ

Writing – original draft: KZS, MYZ

Writing – review & editing: all authors

## Competing interests

The authors declare no competing interests.

**Further information and requests for resources and reagents should be directed to and will be fulfilled by the lead contact, Melody Y. Zeng**

## STAR Methods

### Animals

Wild-type (WT) C57BL/6 mice were originally purchased from the Jackson Laboratory and maintained and expanded in-house by the Zeng laboratory in both germ-free (GF) or specific pathogen-free (SPF) conditions. *Tph1^fl/fl^* mice (C57BL/6 background) was generously provided by Dr. David Artis (Flamar et al., 2020). *Villin^cre^*mice (C57BL/6 background) were purchased from Jackson Laboratory. *Tph1^fl/fl^Villin^cre^*mice were obtained by crossing *Tph1^fl/fl^ with Villin^cre^.* All animal experiments were approved by the Institutional Animal Care and Use Committee at Weill Cornell Medicine.

### Human infant stool specimens

Stool specimens from term infants who were in their first 2 weeks of life, never treated with antibiotics, and admitted to Well Baby Nursery at NewYork Presbyterian-Weill Cornell Medicine were part of a study approved by the NewYork Presbyterian-Weill Cornell Medicine Institutional Review Board.

### CE-TOFMS-based metabolome analysis

Luminal contents from the ileum of SPF WT adult (8 weeks old) and neonatal (2 weeks old) mice were submitted for high-throughput metabolomics as described previously (Kim et al., 2017b). Luminal samples were freeze dried and disrupted by vigorous shaking at 1,500 rpm for 10 min with four 3 mm zirconia beads by Shake Master NEO (Bio Medical Science Inc.). 10 mg (±0.5 mg) fecal samples were homogenized with 500 µl MeOH containing internal standards (20 µM each of methionine sulfone, and D-camphor-10-sulfonic acid (CSA)) and 100 mg of 0.1 mm and four of 3 mm zirconia/silica beads (BioSpec Products). After vigorous shaking (1,500 rpm for 5 min) by Shake Master NEO (Bio Medical Science Inc.), 200 µl of Milli-Q water and 500 µl of chloroform was added and then shaking in a same manner as before. After centrifugation at 4,600*g* for 15 min at 4°C, the supernatant was transferred to a 5 kDa cutoff centrifugal filter tube. The filtrate was centrifugally concentrated at 40°C and reconstituted with 40 µl of Milli-Q water. Ionic metabolites were analyzed using CE-TOFMS in both positive and negative mode. All CE-TOFMS experiments were performed using an Agilent CE capillary electrophoresis system (Agilent Technologies, Inc.). To identify peak annotation and quantification, the obtained data were processed using in-house software (MasterHands).

### Preparation of samples for liquid chromatography/mass spectrometry (LCMS)

Small intestine and colon luminal contents were weighed, placed into 1.5 mL tubes, dissolved in 50% methanol, and vortexed at the highest speed setting for 15 min. Small intestine, colon, and brain tissues were dissolved in 50% methanol and homogenized using an OMNI Bead Rupter 12 (OMNI International). Plasma was mixed with methanol at a 1:1 ratio and vortexed at maximum speed for 15 min. Samples were centrifuged at the maximum speed for 25 min and the supernatant was collected for LCMS. D3-Leucine was added into the solvent at a 1:5 ratio as an internal standard control.

### LCMS detection of 5-HT

The supernatant of tissues was analyzed using an Agilent 1290 LC system coupled to an Agilent 6530 quadrupole time-of-flight (QTOF) mass spectrometer with a 1.7 µm, 2.1 × 100 mm ACQUITY UPLC® BEH C18 column (Agilent). The following solvent system for serotonin detection was used: A: H_2_O with 0.1% formic acid; B: Methanol with 0.1% formic acid. The flow rate was 0.35 mL/min with a column temperature of 40°C. The gradient for HPLC-MS analysis was: 0-4.0 min 99.5% A – 30.0% A, 4.0-4.5 min 30.0% A-2.0%A, 4.5-5.4 min 2.0%A-2.0%A, 5.4-5.6min 2.0%A – 99.5% A. Peaks were assigned by comparison with authentic standards and the absolute analyte concentrations were quantified based on a standard curve derived from a serotonin concentration gradient and associated Peak areas.

### Immunofluorescence staining of 5-HT, MAO-A, and avidin in mouse tissues

Mouse tissues were fixed using 4% paraformaldehyde (PFA, Thermo Fisher) overnight. Tissues were washed with PBS and then placed into 30% sucrose overnight. Tissues were then placed into Tissue-Tek O.C.T. gel (Sakura) and 5 µM sections were placed onto Superfrost Plus microscope slides (Thermo Fisher). Tissue sections were permeabilized with 0.5% TritonX100 for 15 min at room temperature. Antigen retrieval was conducted on tissues using Citrate Buffer, pH 6.0 Antigen Retriever (Sigma) for 30 min at 85°C. After a wash step, sections were blocked with 5% horse serum in PBS for 60 min at room temperature, then incubated with goat-anti-5-HT (1:500, Abcam) or rabbit anti-mouse MAO-A (1:500, Abcam) at 4°C overnight. After a wash step, sections were incubated with donkey anti-goat IgG H&L Alexa Fluor 488 antibody (1:1000, Abcam), anti-rabbit IgG H&L Alexa Fluor 555 antibody, and/or anti-avidin-rhodamine (1:2000, Thermo Fisher) for 60 min, then counterstained with DAPI (1:10000) for 5 min. After staining, coverslips were mounted onto tissue sections using ProLong™ Gold Antifade Mountant (Invitrogen). Images were captured using a Zeiss AxioObserver inverted fluorescence microscope.

### RT-qPCR of gene expression

Total RNA was extracted from cells, colon organoids, or mouse tissue samples after homogenization using TRIzol reagent (Invitrogen) according to the manufacturer’s instructions. RNA (2000 ng per sample) was reverse transcribed into cDNA using a High Capacity cDNA Reverse Transcription kit (Applied Biosystems) according to the manufacturer’s instructions. cDNA was diluted to 20 ng/µL in water before performing qPCR. qPCR was carried out with the CFX384 Real-Time System C1000 Touch Thermal Cycler (Bio-Rad Laboratories) and using the ig™ SYBR Green qPCR 2X Master Mix (Intact Genomics, Inc.) according to the manufacturer’s instructions. Primer sequences used for qPCR are listed in **Table S4**. Relative expression was calculated using the ΔΔCt method, using *B-actin* as the reference gene for all samples.

### Western blot of MAO-A

Tissues were homogenized in RIPA buffer with protease and phosphatase inhibitors, and tissue lysates were collected. Protein concentrations in the lysates were measured using a Pierce™ BCA Protein Assay Kit (Thermo Fisher). Lysates were normalized and then resolved using SDS/PAGE gels (Invitrogen) and transferred onto a PVDF membrane (Bio-Rad Laboratories). Membranes were blocked using 3% non-fat milk for 1 h at room temperature and then probed with rabbit anti-mouse MAO-A antibody (1:500, Abcam) at 4°C overnight. Membrane was probed with goat anti-rabbit-HRP antibody (1:1000, Invitrogen) for 1 h at room temperature. SuperSignal™ West Pico PLUS Chemiluminescence Substrate (Thermo Scientific) was added to the membrane, and the blot was imaged using a ChemiDoc™ XRS+ Molecular Imager (Bio-Rad Laboratories). Beta-actin was used as the loading control. Densitometry analyses of protein bands were performed using ImageJ software.

### Colon organoid culture and maintenance

Colon organoids were kindly given to us by Dr. M. Laura Martin. Fresh tissue biopsy samples were placed in DMEM media (Invitrogen) with GlutaMAX (1×, Invitrogen), 100 U/ml Penicillin-Streptomycin (Gibco) and Primocin 100 µg/ml (InvivoGen). Tissue samples were washed in media two times before being placed in a 10 cm petri dish (Falcon) for mechanical dissection. The dissected tissue was then enzymatically digested with 250 U/ml of collagenase IV (Life Technologies) with 10 µM ROCK inhibitor (Selleck Chemical Inc.) in a 15 ml conical centrifuge tube (Falcon) incubated in a shaker at 37 °C set to 200 rpm. Incubation time of the specimen was dependent on the amount of collected tissue and ranged from 20 to 60 min, until the majority cell clusters were in suspension. After tissue digestion, Advanced DMEM/F12 media (Invitrogen) containing GlutaMAX (1×, Invitrogen), 100 U/ml penicillin-streptomycin (Sigma), and HEPES (10 mM, Gibco) (+++ media) was added to the suspension and the mixture was centrifuged at 326 rcf for 3 min. The pellet was then washed with +++ media. The pellet was resuspended with colon-specific culture media. The final resuspended pellet was combined with Matrigel (Corning) in a 1:2 volume Matrigel, with 5 80 µl droplets pipetted onto each well of a six-well suspension culture plate (GBO). The plate was placed into a cell culture incubator at 37 °C and 5% CO_2_ for 30 min to solidify the droplets before 3ml of colon-specific culture media was added to each well. The culture was maintained with fresh media changed twice a week. Dense cultures with organoids ranging in size from 200 to 500 um were passaged weekly. During passaging, the organoid droplets were mixed with TrypLE Express (Gibco) and placed in a water bath at 37 °C for a maximum of 7 min. The resulting cell clusters and single cells were washed and re-plated, following the protocol listed above. Colon cancer organoids were biobanked using Recovery Cell Culture Freezing Medium (Gibco) at −80 °C and after 24h transferred to liquid nitrogen. Throughout colorectal organoid development and maintenance, cultures were screened for various Mycoplasma strains using the PCR Mycoplasma detection kit (ABM) and confirmed negative before being used for experimental assays.

### Cell line maintenance

HT-29 cells were used in cellular assays. Cells were routinely cultured in McCoy’s 5A (Modified) Medium (Gibco) supplemented with 10% fetal bovine serum (FBS; Gibco) and 1% penicillin-streptomycin in a 37 °C incubator under an atmosphere with 5% CO_2_.

### 16S rRNA sequencing

Bacterial genomic DNA was extracted from fecal pellets and luminal contents using the E.Z.N.A stool DNA kit (Omega Biotek) and DNA amplicons of the V4-V5 region within the 16S rRNA gene was sequenced by the Genomics Core Facility of the WCM Core Laboratories Center. The sequences were curated using Mothur (v.1.35) and sequences were binned into OTUs at >97 % sequence level. LEfSe Linear discriminant analysis (LDA) was performed by the Mothur command using all OTU reads. The OTUs that were present in >0.1 % abundance at maximum were selected and shown in Figure 1B and Table S1. The OTUs shown in the LEfSe panels are those whose abundance was statistically different with a p-value <0.05.

### ELISA of 5-HT

For measurement of 5-HT in tissues, goat anti-5-HT antibody (Abcam, 1:500) was used to coat 96-well ELISA plates (Nunc MaxiStop flat bottom 96-well, Thermo Scientific) overnight at 4°C. Tissues were homogenized in sterile PBS with 1% BSA and separated from debris/mouse cells by removing the pellet after centrifugation at 15,000 rpm for 1 min. 100 µL of supernatant was added to each well overnight at 4°C. After washing, 100 uL of rat anti-5-HT antibody diluted in PBS with 1% BSA (Novus Biologics, 1:1000) was added to each well and incubated at room temperature for 2 h. After washing, goat-anti-rat HRP-conjugated IgG (Bethyl Laboratories, 1:10000) was added to the plate and incubated at room temperature for 1h. After wash, 1-step Ultra TMB ELISA substrate (Thermo Scientific) was added and the plate was incubated for 5 minutes before adding stop solution (Invitrogen). Absorbance was measured at 450 nm using a SpectraMax plate reader.

### Bacterial isolation and Sanger sequencing

Luminal contents from SPF WT neonatal (2 weeks old) mice were collected within an anaerobic chamber and dissolved in PBS. The contents were then dissolved and serial diluted into BHI, Columbia medium supplemented with sheep’s blood, and MRS broth and then plated in their respective agar medium. For isolation of anaerobic bacteria, plates were placed at 37 °C in the anaerobic chamber. For isolation of aerobic bacteria, plates were placed in an incubator at 37 °C. The bacteria were allowed to grow overnight, and single colonies were picked and placed into their respective growth mediums. Bacteria were then grown into log phase and then were used in downstream applications. Bacterial DNA was isolated using the E.Z.N.A Stool DNA kit (Omega Bio-Tek Inc) according to the manufacturer’s instructions. The 16S rRNA gene was PCR amplified using the 8F and 1492R universal primers (Turner et al., 1999). PCR products were submitted to GENEWIZ (Azenta Life Sciences) for Sanger sequencing and the bacterial isolates were identified using Microbial Nucleotide BLAST (NCBI).

### Treatment of HT-29 cells with bacterial isolate supernatant

HT-29 cells were seeded at 10,000 cells/well in a 24-well plate and treated with 22 um-filtered supernatants (equivalent to 10^7^ CFUs) dissolved in McCoy’s 5A (Modified) Medium (Gibco) supplemented with 10% FBS (Gibco) and 1% penicillin-streptomycin in a 37 °C incubator under an atmosphere with 5% CO_2_. After 24 h, the cells were washed and harvested for total RNA isolation using Trizol reagent (Invitrogen) according to the manufacturer’s instructions.

### Colonization of bacterial isolates in GF neonates/dams

AB21 and CD2 are two isolates from SPF P14 mouse small intestine (**Fig. 2H**, **Table S2**). AB21 was grown in BHI liquid growth medium in aerobic conditions. CD2 was grown in Columbia liquid broth medium in an anaerobic chamber at 37°C overnight. The next day, the bacteria were resuspended in sterile PBS and 10^6^ cells were given to GF dams at P2 via oral gavage. After two weeks, neonates were sacrificed for further analyses.

### Isolation of mouse intestinal lamina propria cells

Mouse intestines were removed, cleaned from remaining fat tissue, and washed in ice-cold PBS (Corning). Intestines were opened longitudinally, washed in ice-cold PBS, and cut into small pieces (∼2 mm). Dissociation of epithelial cells was performed by incubation on a shaker in Hanks’ balanced salt solution (Sigma-Aldrich) containing 5 mM EDTA (Thermo Fisher Scientific), 1 mM dithiothreitol (DTT) (Sigma-Aldrich), and 2% heat-inactivated fetal bovine serum (FBS) for 15 min at 37°C with agitation. The dissociated cells were filtered through a 70 µm cell strainer and washed one time prior to enzymatic digestion in buffer containing dispase (0.4 U/ml; Thermo Fisher Scientific), collagenase III (1 mg/ml; Worthington), and deoxyribonuclease (DNase) I (20 µg/ml; Sigma-Aldrich) in RPMI medium (Sigma Aldrich) containing 10% FBS for 25-30 min at 37°C. Leukocytes were further enriched by a 40/75% Percoll gradient centrifugation (Cytiva) at 2000 rpm for 20 min with no brake.

### Isolation of splenic CD3+ T cells and in vitro treatment/culture

SPF WT B6 mice were sacrificed and the spleen was harvested. Splenocytes were processed into a single cell suspension and red blood cells were lysed using ACK lysis buffer and rinsed with PBS. Cells were incubated with anti-CD3-APC antibody (Invitrogen) for 20 min on ice then rinsed with PBS. Cells were then incubated with anti-APC MACS Microbeads (Miltenyi Biotec) for 10 min on ice then rinsed with PBS. Cells were passed through a MACS bead column according to the manufacturer’s directions. Remaining CD3+ T cells were cultured for 24-48 h with 0-40 µM 5-HT in RPMI medium supplemented with 10% FBS, anti-mouse-CD3e antibody (1µg/mL, Invitrogen), and anti-mouse-CD28 antibody (0.5 µg/mL, Invitrogen). Cells were then analyzed using flow cytometry (methods below).

### Oral gavage of GF neonatal mice with PBS or 5-HT (schematic in Fig. 4F)

GF WT neonatal mice were given 50 µL of 5-HT (12.5 µg/mL) or vehicle (PBS) at P7 and P8 via oral gavage. The mice were then sacrificed at P14 and SI and colon tissues were collected for further analysis of lamina propria immune cells. Cells were analyzed using flow cytometry (methods below).

### Oral tolerance to commensal bacteria (schematic in Fig. 5D)

GF WT neonatal mice were given 50 µL of 5-HT (12.5 µg/mL) or vehicle (PBS) at P7 and P8 via oral gavage. When the neonates were P14, their dams were given 200 µL of SI and colon luminal contents (dissolved in PBS) from P14 SPF WT mice. After 2 weeks, to allow bacterial colonization, the mice were sacrificed and SI and colon were collected for lamina propria immune cell analysis by flow cytometry. SI luminal contents and fecal pellets were also collected for 16S rRNA sequencing.

### Oral tolerance to OVA (schematic in Fig. 5A)

GF WT neonatal mice were given 50 µL of 5-HT (12.5 µg/mL) or vehicle (PBS) at P7 and P8 via oral gavage. The dams were then oral gavaged with 200 µg ovalbumin (OVA, Sigma Aldrich) dissolved in 200 µL PBS when neonates were P12, P15, P18, and P21, allowing the neonatal mice to be sensitized to OVA via the dam’s breastmilk. At P28, the mice were weaned from the dams. When mice were 5 weeks old, they were challenged with OVA dissolved in Freund’s complete adjuvant via subcutaneous (s.c.) footpad injection. When mice were 7 weeks old, they were re-challenged with OVA dissolved in Freund’s incomplete adjuvant via s.c. footpad injection. 3 days after the re-challenge, mice were sacrificed and their blood and spleen was collected for further analysis of OVA-specific antibodies and immune cell characterization via flow cytometry, respectively.

### ELISA of OVA-specific IgG, IgE, and IgM

For measurement of OVA-specific Igs in plasma, OVA (2 µg/mL, Sigma Aldrich) was used to coat 96-well ELISA plates (Nunc MaxiStop flat bottom 96-well, Thermo Scientific) overnight at 4°C. Plasma was diluted (1:10 for detection of OVA-specific IgG and IgM; 1:5 for detection of OVA-specific IgE) in sterile PBS with 1% BSA and added to each well at 4°C overnight. After washing, HRP-conjugated anti-mouse-IgG, HRP-conjugated anti-mouse IgE, or HRP-conjugated anti-mouse IgM (all diluted at 1:10000) were added to the plate. After a final wash, 1-step Ultra TMB ELISA substrate (Thermo Scientific) was added and the plate was incubated for 5 minutes before adding stop solution (Invitrogen). Absorbance was measured at 450 nm using a SpectraMax plate reader. HRP-conjugated antibodies and HRP substrates were purchased from Bethyl Laboratories.

### Flow cytometry

Lamina propria cells from mouse intestines were isolated as previously described, followed by stimulation for 4 h ex vivo with phorbol 12-myristate 13-acetate (PMA; 100 ng/ml; Sigma-Aldrich) and ionomycin (1 µg/ml; Sigma-Aldrich) in a 37°C incubator (5% CO_2_). Golgi plug was added to cells after 2 h of stimulation (1:1000, BD Biosciences). Cells were washed with cold PBS, blocked with Fc block for 15 min, resuspended in FACS buffer (PBS with 1% BSA) and stained for surface markers and viability for 20 min (all antibodies used are listed in **Table S5**). For intracellular staining, cells were fixed and permeabilized using the eBioscience Foxp3/Transcription Factor Staining Buffer Set according to the manufacturer’s protocol (eBioscience). Stained cells were then washed with FACS buffer once, and analyzed by Cytek Aurora (Cytek). Data were analyzed using Flow Jo (Becton Dickinson).

### Seahorse mitochondrial stress test

SPF WT B6 adult mice were sacrificed and the spleen was harvested. Splenocytes were processed into a single cell suspension and red blood cells were lysed using ACK lysis buffer and rinsed with PBS. Cells were incubated with anti-CD3-APC antibody (Invitrogen) for 20 min on ice then rinsed with PBS. Cells were then incubated with anti-APC MACS Microbeads (Miltenyi Biotec) for 10 min on ice then rinsed with PBS. Cells were passed through a MACS bead column (Miltenyi Biotec) according to the manufacturer’s directions. Remaining CD3+ T cells were seeded at 300,000 cells/well onto a tissue culture microplate (Seahorse XFp FluxPak, Agilent) pre-treated with Cell-Tak Solution (Corning) and cultured for 3 h with 0-20 µM 5-HT in Seahorse XF RPMI media (Agilent Technologies) supplemented with 25mM glucose (Agilent Technologies), 2mM L-glutamine (Agilent Technologies), and 1mM pyruvate (Agilent Technologies). After 3 h, a Seahorse Mito Stress Test Assay was performed using a Seahorse XFp Mito Stress Kit (Agilent Technologies) according to the manufacturer’s instructions and as previously described (Van der Windt et al., 2016). Oxygen consumption rate was measured using a Seahorse XFp Analyzer (Agilent Technologies).

### Statistical analyses

Statistical analyses were performed using GraphPad Prism 7.0 (GraphPad Software). Differences between two groups were evaluated using Student’s t test (parametric) or Mann-Whitney U test (non-parametric). For multiple comparisons, one-way ANOVA (parametric) or Kruskal-Wallis test (non-parametric) were used. Differences with p-values < 0.05 were considered significant.

