## Supplemental Information for "Gut bacteria-derived serotonin promotes immune tolerance in early life"

### Supplemental Figure 1

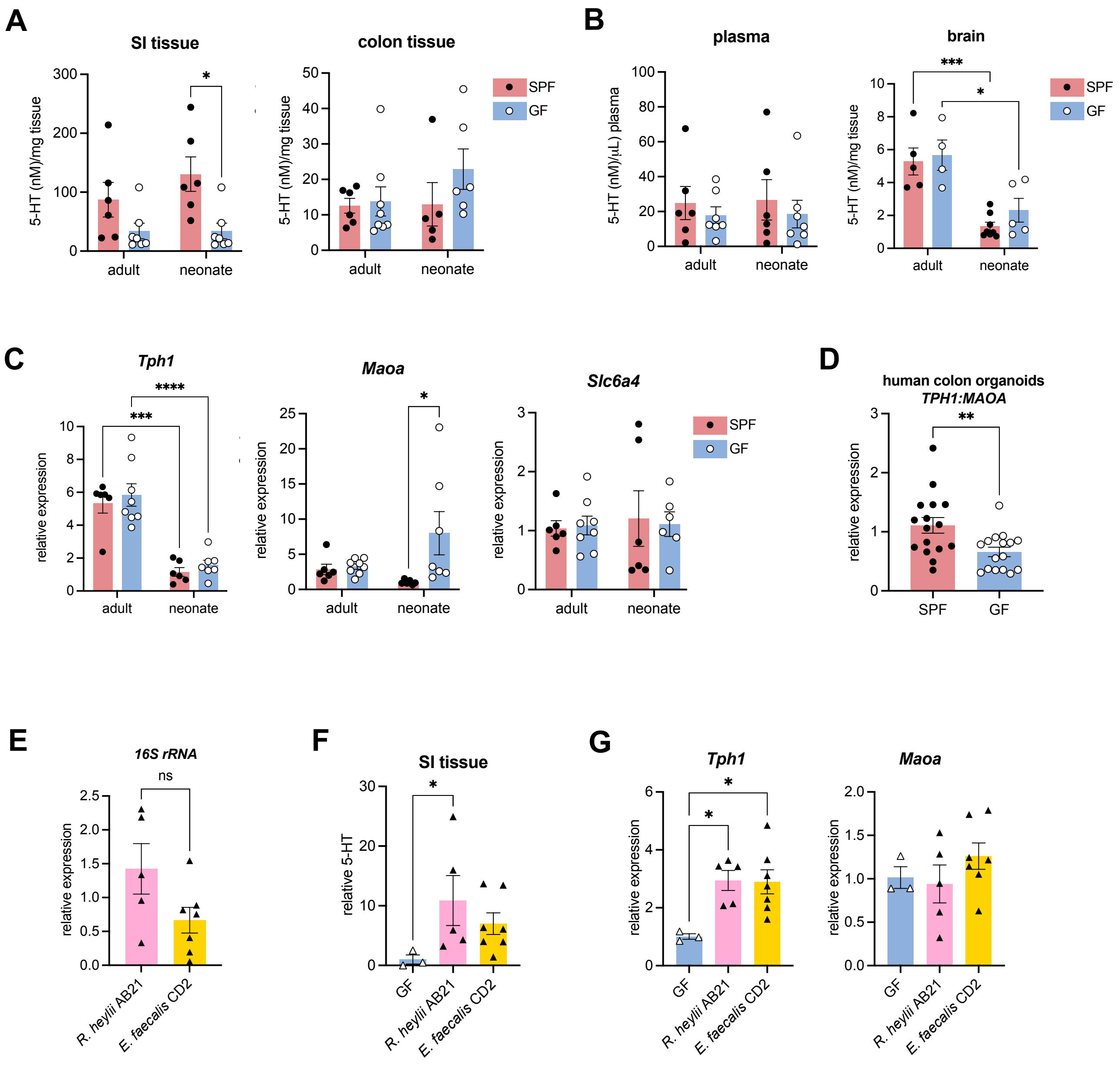

**Fig. S1, related to Fig. 2. 5-HT biosynthesis in the neonatal intestine is driven by the gut microbiome.** (**A-B**) LCMS analysis of 5-HT in (A) SI and colon tissues, and (**B**) plasma and brain of SPF/GF adult (> 8 weeks) and neonatal (2-week-old) mice. (**C**) RT-qPCR analysis of *Tph1, Maoa,* and *Slc6a4* gene expression in the colons of SPF/GF adult and neonatal mice. (**D**) RT-qPCR analysis of *TPH1:MAOA* expression ratio in human colon organoids treated with SI luminal contents from SPF and GF neonatal mice. (**E-G**) (**E**) *16S rRNA* gene expression in feces, (**F**) ELISA analysis of 5-HT in SI tissues, and (**G**) RT-qPCR analysis of *Tph1* and *Maoa* in SI of GF neonatal mice colonized with bacterial isolates AB21 or CD2 after 2 weeks. *$P$<0.05, **$P$<0.01, ****$P$<0.000. Adult mice age > 8 weeks, neonatal mice age = 2 weeks.

### Supplemental Figure 2

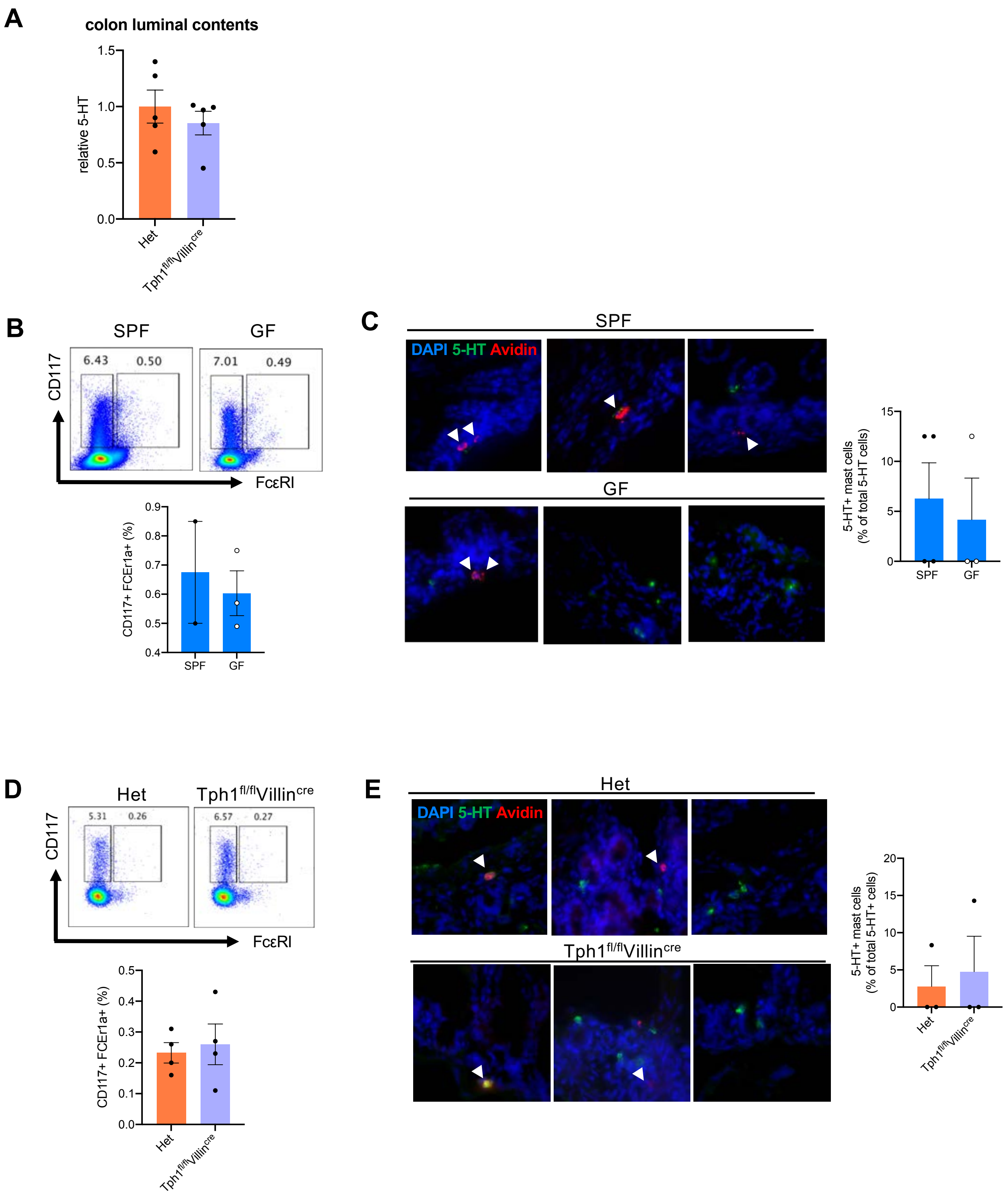

**Fig. S2, related to Fig. 3. Mast cells are not major 5-HT producers in the neonatal mouse gut.** (**A**) 5-HT concentrations in colon luminal contents of *Tph1*$^{fl/fl}$*Villin*$^{cre}$ neonatal (2-week-old) mice or heterozygous (Het) littermates measured by ELISA. (**B**) Representative flow plots and quantification of mast cell populations in SI lamina propria of SPF and GF neonatal mice. (**C**) Immunofluorescence staining and quantification of 5-HT positive mast cells in SI of SPF and GF neonatal mice. (**D**) Representative flow plots and quantification of mast cell populations in SI lamina propria of *Tph1*$^{fl/fl}$*Villin*$^{cre}$ neonatal mice or Het littermates. (**E**) Immunofluorescence staining and quantification of 5-HT positive mast cells in SI of *Tph1*$^{fl/fl}$*Villin*$^{cre}$ neonatal mice or Het littermates.

### Supplemental Figure 3

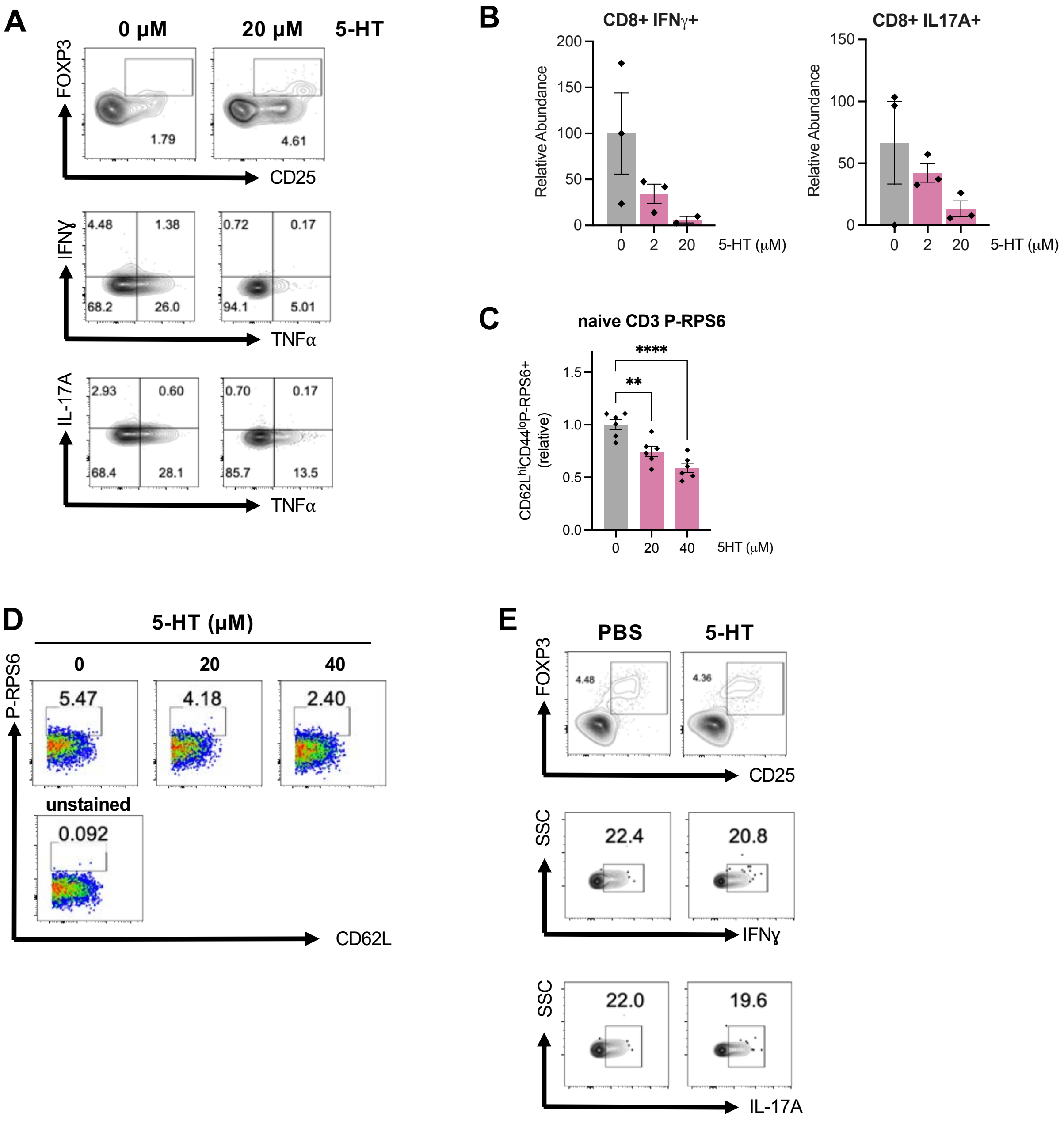

**Fig. S3, related to Fig. 4. 5-HT promotes Treg development *in vitro* and *in vivo* in the neonatal SI.** (**A**) Representative flow plots of mouse splenic T cells stimulated with 5-HT *in vitro*. (**B**) Flow cytometry analysis of mouse splenic CD8+ T cells stimulated with 5-HT *in vitro* for 48 h. (**C**) Flow cytometry analysis of mouse splenic naïve CD3+ P-RPS6+ T cells stimulated with 5-HT *in vitro* for 24 h. (**D**) Representative flow plots of mouse naïve CD4+ P-RPS6+ T cells treated with or without 5-HT. (**E**) Representative flow plots of lamina propria immune cells from colon of GF neonatal mice treated with PBS or 5-HT.

### Supplemental Figure 4

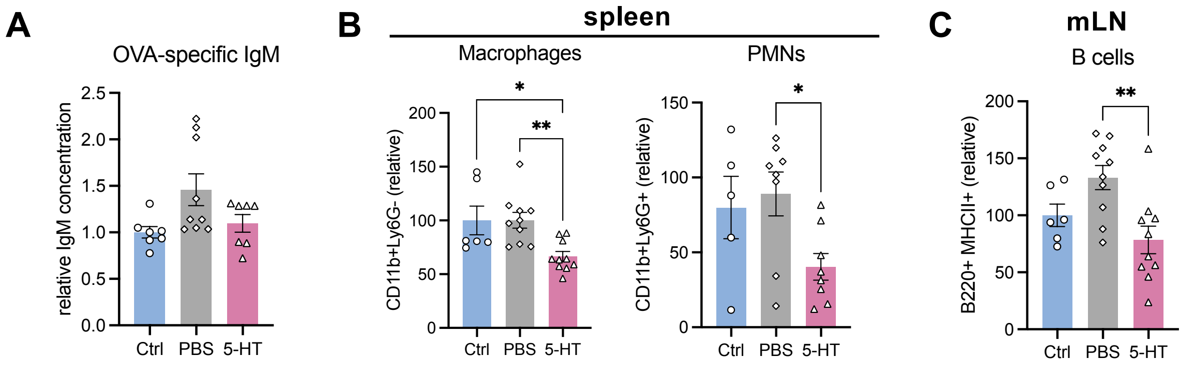

**Fig. S4, related to Fig. 5. 5-HT in the neonatal gut promotes long-term immune tolerance to oral antigens.** (**A**) OVA-specific IgM in the plasma of GF mice treated with PBS or 5-HT and challenged with OVA. (**B-C**) Flow cytometry analysis of immune cells isolated from the (**B**) spleen and (**C**) mLN of GF mice treated with PBS or 5-HT and challenged with OVA. Neonatal mice age = 2 weeks. *$P$<0.05, **$P$<0.01.

### Supplemental Figure 5

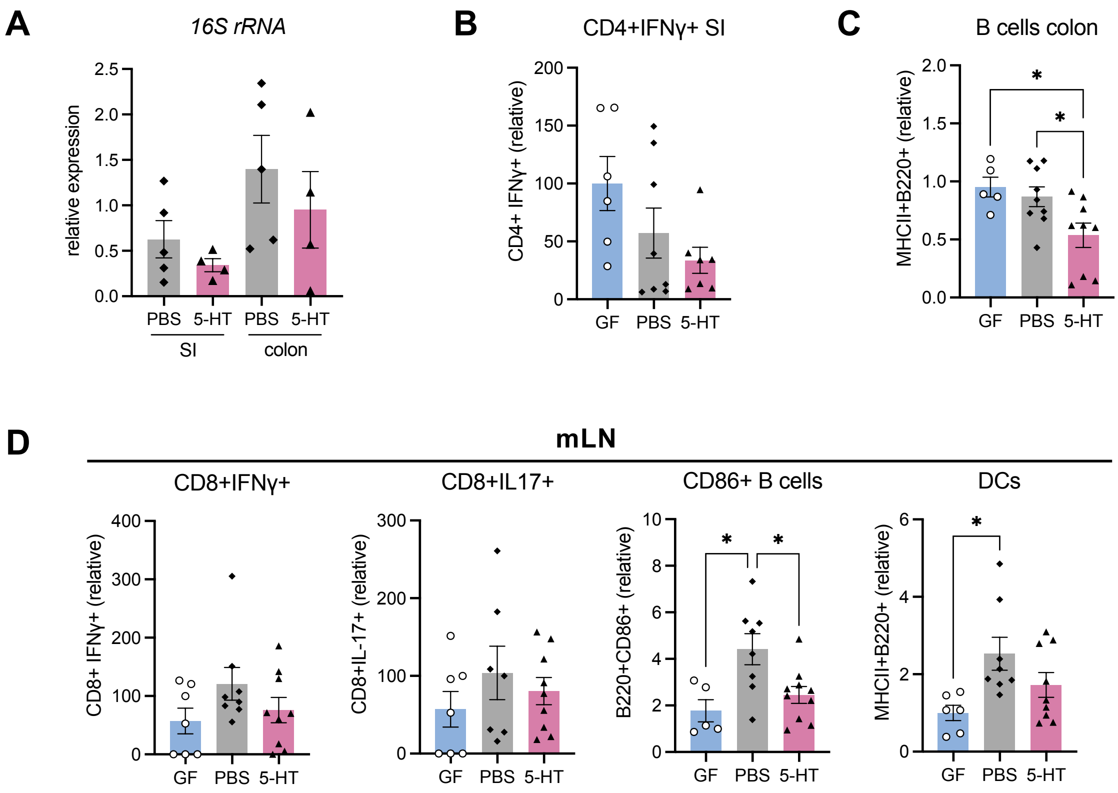

**Fig. S5, related to Fig. 6. 5-HT in the neonatal gut promotes immune tolerance to gut commensal bacteria.** (**A**) 16S rRNA expression in SI and colon luminal contents of GF mice treated with PBS or 5-HT and then colonized with commensal bacteria for 2 weeks. (**B-C**) Flow cytometry analysis of immune cells isolated from the (**B**) SI, (**C**) colon, and (**D**) mLN of GF neonatal mice treated with PBS or 5-HT, followed by colonization of the dam with gut commensal bacteria isolated from P14 SPF mice for 2 weeks. All mice were housed in the same cage. Neonatal mice age = 2 weeks. *$P<0.05$, **$P<0.01$.

**Supplemental Figure 6**

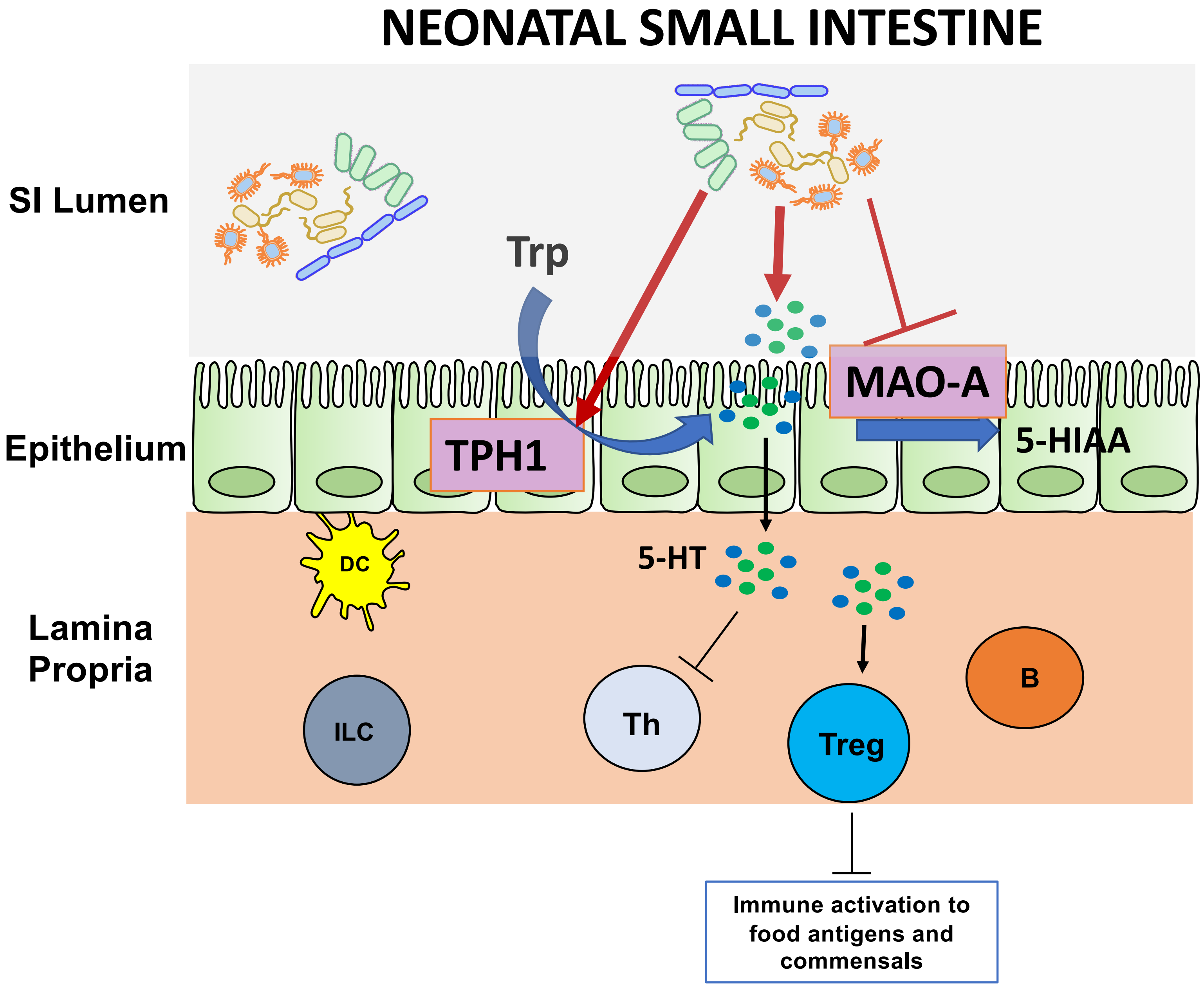

**Fig. S6. Model of gut-microbiota dependent 5-HT synthesis and its subsequent immunomodulatory effects in the neonatal intestine.** In the neonatal intestine, 5-HT is produced directly by bacteria and indirectly through the increase of TPH1 and suppression of MAO-A in intestinal enterochromaffin cells. The gut-derived 5-HT can induce Tregs and suppress immune activation to food antigens and commensals.

**Table S1. OTU identities with highest homology of bacteria from LDA analysis, related to Fig. 1B.**

| OTUs | Bacterium identity (highest homology) | GenBank ID | 16S identity match (%) |
|---|---|---|---|
| Otu000001_Lactobacillus | *Lactobacillus murinus* strain V10 | CP040852.1 | 100 |
| Otu000023_Staphylococcus | *Staphylococcus saprophyticus* strain ARSS3 | MT317182.1 | 100 |
| Otu000157_Bacilli_unclassified | *Staphylococcus sp.* 219 | GQ222242.1 | 97.326 |
| Otu000199_Bacilli_unclassified | *Lactobacillus murinus* | EF187260.2 | 95.989 |
| Otu000147_Lactobacillales_unclassified | *Enterococcus faecium* strain Mise766 | MN994406.1 | 96.524 |
| Otu000156_Bacilli_unclassified | *Lactobacillus murinus* | EF187260.2 | 96.791 |
| Otu000842_Lactobacillales_unclassified | *Lactobacillus murinus* strain V10 | CP040852.1 | 96.524 |
| Otu000545_Streptococcus | *Streptococcus salivarius* strain D32-2 | MT275480.1 | 100 |
| Otu000169_Lactobacillus | *Lactobacillus murinus* strain V10 | CP040852.1 | 96.791 |
| Otu000140_Coriobacteriaceae_unclassified | *Adlercreutzia sp.* strain D16-63 | MK287694.1 | 91.622 |
| Otu000134_Coriobacteriaceae_unclassified | *Enterorhabdus sp.* strain 21x | MK287690.1 | 93.011 |
| Otu000094_Firmicutes_unclassified | *Lactobacillus johnsonii* strain TP6.10 | MF193450.1 | 94.309 |
| Otu000109_Coriobacteriaceae_unclassified | *Enterorhabdus sp.* strain 21x | MK287690.1 | 92.992 |
| Otu000090_Lachnospiraceae_unclassified | *Murimonas sp.* strain 102X | MK287626.1 | 91.176 |
| Otu000111_Coriobacteriaceae_unclassified | *Enterorhabdus sp.* strain 21x | MK287690.1 | 93.531 |
| Otu000172_Firmicutes_unclassified | *Lactobacillus gasseri* strain A4 | MK982449.1 | 94.737 |
| Otu000104_Ruminococcaceae_unclassified | *Anaerotruncus sp.* strain 0.1XD8_49 | MN081629.1 | 99.731 |
| Otu000077_Ruminococcaceae_unclassified | *Monoglobus pectinilyticus* strain ASD1037 | MK615117.1 | 94.385 |
| Otu000070_Lactobacillales_unclassified | *Lactobacillus gasseri* clone WWC_C2MKM009 | GU417794.1 | 95.135 |
| Otu000085_Firmicutes_unclassified | *Rummeliibacillus suwonensis* strain JA28 | MT269581.1 | 94.086 |
| Otu000075_Lactobacillales_unclassified | *Lactobacillus gasseri* clone WWC_C2MKM060 | GU417806.1 | 97.059 |
| Otu000051_Clostridiales_unclassified | *Clostridiales bacterium* oral taxon F32 strain VO026 | HM099644.1 | 95.979 |
| Otu000047_Lachnospiraceae_unclassified | *Lachnospiraceae bacterium* DW44 | KX009924.1 | 94.595 |
| Otu000044_Lachnospiraceae_unclassified | *Dorea formicigenerans* strain DSM 105840 | MN537498.1 | 96.515 |

| | | | |
|---|---|---|---|
| Otu000039_Clostridiales_unclassified | Uncultured bacterium clone LCM019 | JF733451.1 | 99.196 |
| Otu000041_Porphyromonadaceae_unclassified | *Bacteroidales bacterium* strain NM04_E33 | MK929052.1 | 92.493 |
| Otu000061_Firmicutes_unclassified | *Lactobacillus acidophilus* strain Bonyadi | KF056894.1 | 89.84 |
| Otu000034_Porphyromonadaceae_unclassified | *Muribaculum sp.* strain 26x | MK287695.1 | 100 |
| Otu000028_Porphyromonadaceae_unclassified | *Muribaculum sp.* TLL-A4 | CP039393.1 | 95.71 |
| Otu000031_Porphyromonadaceae_unclassified | *Porphyromonas sp.* k30 | KT263531.1 | 93.617 |
| Otu000022_Bacteroidales_unclassified | *Bacteroidales bacterium* strain 32x | MK287702.1 | 92.493 |
| Otu000017_Porphyromonadaceae_unclassified | *Bacteroidales bacterium* | LC333640.1 | 94.841 |
| Otu000010_Lachnospiraceae_unclassified | *Lachnospiraceae bacterium* DW44 | KX009924.1 | 100 |
| Otu000008_Turicibacter | *Turicibacter sp.* strain TS-3 | MK287738.1 | 100 |
| Otu000003_Clostridiales_unclassified | *Candidatus Arthromitus sp.* SFB-mouse-NL | CP008713.1 | 100 |
| Otu000002_Lactobacillus | *Lactobacillus johnsonii* strain G2A | CP040854.1 | 100 |

**Table S2. Bacterial isolates identified as up-regulators of *Tph1* or *Maoa*, related to Fig. 2D.**

| **Upregulators of TPH1** | | |
|---|---|---|
| **Isolate** | **Identity** | **Relative expression of *TPH1*:*MAOA*** |
| AC15 | *Enteroccocus gallinarum* | 14.1 |
| AC17 | *Enteroccocus gallinarum* | 598.6 |
| AB21 | *Rodentibacter heylii* | 12,690.2 |
| AB22 | *Rodentibacter heylii* | 2,393.7 |
| AB23 | *Staphylococcus xylosus* | 8,438.1 |
| AB26 | *Staphylococcus xylosus* | 19,483.3 |
| AB2 | *Rodentibacter heylii* | 7,601.9 |
| BB1 | *Enteroccocus gallinarum* | 13.7 |
| BB2 | *Enteroccocus gallinarum* | 12.9 |
| CB1 | undetermined | 4.1 |
| **Upregulators of MAO-A** | | |
| **Isolate** | **Identity** | **Relative expression of *TPH1*:*MAOA*** |
| AF1 | *Pasteurella caecimuris* | 0.003 |
| CD2 | *Enteroccocus faecalis* | 0.016 |
| CD3 | *Enteroccocus faecalis* | 0.043 |

| | | |
|---|---|---|
| F2 | *Streptococcus acidominimus* | 0.04 |

**Table S3. Bacterial isolates from P14 mouse SI capable of producing 5-HT, related to Fig. 3D.**

| **5-HT producers** | |
|---|---|
| **Isolate** | **Identity** |
| AB6 | *Enterococcus gallinarum* |
| AB7 | *Enterococcus gallinarum* |
| AB8 | *Rodentibacter heylii* |
| AB9 | *Enterococcus gallinarum* |
| AB13 | *Enterococcus gallinarum* |
| AB21 | *Rodentibacter heylii* |
| AB22 | *Rodentibacter heylii* |
| AB24 | *Rodentibacter heylii* |
| AB25 | *Rodentibacter heylii* |
| AB26 | *Staphylococcus xylosus* |
| BB1 | *Enteroccocus gallinarum* |
| BB2 | *Enteroccocus gallinarum* |
| BB3 | *Streptococcus acidominimus* |
| B21 | *Streptococcus acidominimus* |
| B23 | *Streptococcus acidominimus* |
| AC20 | *Enteroccocus gallinarum* |
| CC2 | *Enterococcus faecalis* |
| M1 | *Lactobacillus johnsonii* |

**Table S4. Primer sequences for RT-qPCR analyses**

| **Primers for RT-qPCR and sequencing** | | | |
|---|---|---|---|
| **Gene** | **Forward primer** | **Reverse primer** | **Ref.** |
| Mouse *B-actin* | AAGGCCAACCGTGAAAAGAT | GTGGTACGACCAGAGGCATAC | (Hohenstein et al., 2008) |

| | | | |
|---|---|---|---|
| Mouse *Tph1* | AACAAAGACCATTCCTCCGAAAG | TGTAACAGGCTCACATGATTCTC | (De Vadder et al., 2018) |
| Mouse *Maoa* | GCCCAGTATCACAGGCCAC | GTCCCACATAAGCTCCACCA | (Libert et al., 2011) |
| Mouse *Slc6a4* | CTCCGCAGTTCCCAGTACAAG | CACGGCATAGCCAATGACAGA | (Du et al., 2019) |
| Human *B-ACTIN* | GCAAGCAGGACTATGACGAG | CAAATAAAGCCATGCCAATC | (Zeng et al., 2018) |
| Human *GAPDH* | ACAACTTTGGTATCGTGGAAGG | GCCATCACGCCACAGTTTC | (Lee et al., 2019) |
| Human *TPH1* | ACGTCGAAAGTATTTTGCGGA | ACGGTTCCCCAGGTCTTAATC | (Guenin-Macé et al., 2020) |
| Human *MAOA* | GAATCAAGAGAAGGCGAGTATCG | GGCAGCAGATAGTCCTGAAATG | (Zhang et al., 2019) |
| Human *SLC6A4* | ACGGAGTTCTACAGAAGGTTGT | ATAGAGTGCCGTGTGTCATCT | (Wang et al., 2021) |
| Mouse *Il33* | GCTGCGTCTGTTGACACATT | CACCTGGTCTTGCTCTTGGT | (Mahlakõiv et |

| | | | |
|---|---|---|---|
| | | | al., 2019) |
| *16S rRNA* (V3-V4 region for qPCR) | CCTACGGGTGGCTGCAG | GACTACTAGGGTATCTAATCC | (Yang et al., 2018) |
| *16S rRNA* (entire region for Sanger sequencing, 8F, 1492R) | AGAGTTTGATCCTGGCTCAG | GGTTACCTTGTTACGACTT | (Turner et al., 1999) |

**Table S5. Antibodies used for western blot, immunofluorescence staining, and flow cytometry analyses.**

| Antibodies | Clone | Source | Identifier |
|---|---|---|---|
| Donkey Anti-Goat IgG H&L (Alexa Fluor® 488) | Polyclonal | Abcam | ab150129 |
| Alexa Fluor-700 anti-mouse CD45 | 30-F11 | BioLegend | 103128 |
| Brilliant Violet 711 anti-mouse CD3e | 145-2C11 | BioLegend | 100349 |
| Brilliant Violet 510 anti-mouse CD4 | RM4-4 | BioLegend | 116025 |
| PerCP-eFluor 710 anti-mouse CD8a | 53-6.7 | Invitrogen | 46-0081-80 |
| PE anti-mouse CD25 | PC61 | BioLegend | 102007 |
| Brilliant Violet 650 anti-mouse IFN-γ | XMG1.2 | BioLegend | 505832 |
| PE-Cyanine5.5 anti-mouse/rat FOXP3 | FJK-16s | Invitrogen | 35-5773-82 |
| PE/Cyanine7 anti-mouse TNF-α | MP6-XT22 | BioLegend | 506323 |
| PE-Cyanine7 anti-mouse Ly-6G (Gr1) | RB6-8C5 | eBioscience | 25-5931-82 |
| PE anti-mouse/human CD11b | M1/70 | BioLegend | 101207 |
| FITC anti-mouse CD11c | N418 | BioLegend | 117305 |
| Pacific Blue anti-mouse/human CD45R/B220 | RA3-6B2 | BioLegend | 103227 |
| BUV395 anti-mouse CD86 | GL1 | BD Biosciences | 564199 |
| Brilliant Violet anti-mouse IL-17A | TC11-18H10.1 | Biolegend | 506926 |
| PE/Cyanine7 anti-mouse CD117 | 2B8 | Biolegend | 105813 |
| PerCP-Cyanine 5.5 anti-mouse FCER1a | Mar-1 | Biolegend | 134318 |
| Brilliant Violet 650 anti-mouse MHCII (I-A/I-E) | M5/114.15.2 | Biolegend | 107641 |
| PE/Cyanine7 anti-mouse IL-4 | 11b11 | Invitrogen | 25-7041-80 |
| PerCP anti-mouse CD62L | MEL-17 | BioLegend | 104429 |
| Pacific Blue anti-mouse/human CD44 | IM7 | BioLegend | 103019 |
| PE anti-mouse/human phosphor-RPS6 | A17020B | BioLegend | 608603 |
| Rabbit anti-mouse/rat/human MAO-A | EPR7101 | Abcam | ab126751 |
| Alexa Fluor 555 Goat anti-rabbit IgG | polyclonal | Invitrogen | A-21429 |
| Goat anti-serotonin | polyclonal | Abcam | ab66047 |
| Rat anti-serotonin | YC5/45 | Novus Biologicals | NB65037 |

| | | | |
|---|---|---|---|
| Goat anti-rat IgG HRP | polyclonal | Invitrogen | 31470 |
